# Dual targeting of *Saccharomyces cerevisiae* Pso2 to mitochondria and the nucleus, and its functional relevance in the repair of DNA interstrand crosslinks

**DOI:** 10.1101/2022.03.14.484363

**Authors:** Shravanahalli C. Somashekara, Kalappa Muniyappa

## Abstract

Repair of DNA interstrand crosslinks (ICLs) involves a functional interplay among different DNA surveillance and repair pathways. Previous work has shown that ICL-inducing agents cause damage to *Saccharomyces cerevisiae* nuclear and mitochondrial DNA (mtDNA), and its *pso2*/*snm1* mutants exhibit a petite phenotype followed by loss of mtDNA integrity and copy number. Complex as it is, the cause and underlying molecular mechanisms remains elusive. Here, by combining a wide range of approaches with *in vitro* and *in vivo* analyses, we assessed the subcellular localization and function of Pso2. We found evidence that the nuclear-encoded Pso2 contains one mitochondrial targeting sequence (MTS) and two nuclear localization signals (NLS1 and NLS2), although NLS1 resides within the MTS. Further analysis revealed that Pso2 is a dual-localized ICL repair protein; it can be imported into both nucleus and mitochondria, and that genotoxic agents enhance its abundance in the latter. While MTS is essential for mitochondrial Pso2 import, either NLS1 or NLS2 is sufficient for its nuclear import; this implies that the two NLS motifs are functionally redundant. Ablation of MTS abrogated mitochondrial Pso2 import, and concomitantly, raised its levels in the nucleus. Strikingly, mutational disruption of both NLS motifs blocked the nuclear Pso2 import; at the same time, they enhanced its translocation into the mitochondria, consistent with the notion that the relationship between MTS and NLS motifs is competitive. However, the nuclease activity of import-deficient species of Pso2 was not impaired. The potential relevance of dual-targeting of Pso2 into two DNA-bearing organelles is discussed.

## INTRODUCTION

Numerous studies have demonstrated that interstrand DNA crosslinks (ICLs) are extremely toxic to cells (from bacteria to humans) owing to their capacity to physically block unwinding of duplex DNA during cellular processes such as DNA replication, transcription, repair, and recombination: this topic has been extensively reviewed in the literature (Clauson *et al*. 2013; Baddock *et al*. 2020; Kuhbacher and Duxin, 2020; Semlow and Walter, 2021). The crosslinks (involving various nucleobases) can form between complementary strands of DNA, within the same DNA strand, or between DNA and protein (Baddock *et al*. 2020; Semlow and Walter, 2021). These are induced by a plethora of endogenous and exogenous oxidative/nitrosative sources under physiologically relevant conditions, chemical carcinogens and platinum-based anti-cancer drugs (Garaycoechea and Patel, 2014; Rozelle *et al*. 2021; Semlow and Walter, 2021). Indeed, while a single ICL is sufficient to kill repair-deficient, rapidly-dividing bacteria or yeast (Magana-Schwencke*et al*. 1982), analogous experiments with mammalian cells indicate that 20 unrepaired ICLs are detrimental to the survival of these cells (Lawley and Phillips, 1996; Liu and Wang, 2013). Consequently, platinum-based compounds (e.g., cisplatin, carboplatin and oxaliplatin) are currently being used to cure or control various cancers (Deans and West, 2011; Semlow and Walter, 2021).

Mechanistically, ICL repair can be grouped into three categories; transcription-dependent, replication-dependent and transcription- and replication independent pathways (Muniandy *et al*. 2010; Baddock *et al*. 2020; Semlow and Walter, 2021). The bulky ICLs are recognized and removed in the G0/G1 phase of the cell cycle, but those that do not cause significant distortion in the double helix are excised through the replication-coupled pathway in the S phase (Kuhbacher and Duxin, 2020; Semlow and Walter, 2021). Furthermore, ICL repair takes place in non-replicating cells, notably in post-mitotic cells such as neurons and quiescent stem cells (Datta and Brosh, 2019; Baddock *et al*. 2020). The accumulated data indicate that the repair of ICLs requires the complex coordination among multiple nucleases such as XPF-ERCC1, FAN1, MUS81, and SNM1A that belong to different DNA repair pathways (Semlow and Walter, 2021). In line with this, studies have demonstrated that protein factors belonging to three major DNA pathways, namely, nucleotide excision repair (NER), post-replication repair, and homologous recombination-based DNA repair (HR) play a pivotal role in ICL repair in *Saccharomyces cerevisiae* (Williams *et al*. 2013; Lehoczký *et al*. 2007). Parenthetically, growing evidence suggests that ICL repair in vertebrates is more complex than in *S. cerevisiae*. In support of this idea, several enzymes and protein factors involved in base excision repair (BER), Fanconi anaemia (FA), HR, transcription-coupled nucleotide excision repair (TC-NER), double-strand break repair and translesion synthesis are crucial for the repair of ICLs in human cells (Chatterjee and Walker, 2017; Slyskova *et al*. 2018; Semlow and Walter, 2021). Although the underlying mechanism(s) is not fully understood, these DNA repair pathways are thought to function cohesively to orchestrate ICL repair in vertebrate cells.

Indeed, a significant part of our understanding of ICL repair originates from studies on the sensitivity of *S. cerevisiae* strains to DNA crosslinking agents. In particular, Moustacchi and colleagues isolated *S. cerevisiae* mutant strains that were hypersensitive to low doses of various ICL-inducing agents, but not to other types of DNA damaging agents (Henriques and Moustacchi, 1980; Ruhland *et al*. 1981a; 1981b; Henriques *et al*. 1997; Brendel *et al*. 2003). Systematic mutant screens identified ten *PSO* genes, named *PSO1* through *PSO10*. Of these, *PSO2*/*SNM1* (*PSO2*, sensitivity to PSOralen*; SNM1*, Sensitivity to Nitrogen Mustard) has been previously suggested to play a crucial role in the processing of ICL-associated DNA double-strand breaks (Li and Moses, 2003; Barber *et al*. 2005; Dudas *et al*. 2007). Following this discovery, it was demonstrated that the components of the Fanconi-like pathway act redundantly with the Pso2-controlled pathway (Barber *et al*. 2005; Ward *et al*. 2012). The functional homolog of *S. cerevisiae* Pso2 (hereafter referred to as Pso2) in vertebrates is *SNM1A*, as the latter can rescue the *S. cerevisiae pso2Δ* mutants from cisplatin-induced cell death (Hazrati *et al*. 2008). The human *SNM1B* (also called Apollo or DCLRE1B) is one of the three orthologues of the *S. cerevisiae PSO2* gene: its deficiency potentiates genomic instability due to telomere dysfunction and checkpoint deregulation (Demuth *et al*. 2004; van Overbeek and de Lange, 2006; Bae *et al*. 2008). While *SNMA1* and *SNMB1* are redundant in ICL repair, *SNM1A* knockout mice show decreased lifespan due to accelerated tumorigenesis and susceptibility to infection (Dronkert *et al*. 2000; Ahkter *et al*. 2005); the *SNM1B^−/−^* mice exhibit perinatal lethality (Akhter *et al*. 2010). In light of this, *SNM1A* has emerged as a promising drug target for cancer treatment (Baddock *et al*. 2020; Buzon *et al*. 2021).

From a structural perspective, Pso2 is a member of the superfamily of highly-conserved metallo-β-lactamase/β-CASP fold containing nucleic acid processing enzymes with roles in ICL and DSB repair as well as regulation of cell cycle checkpoints (Aravind *et al*. 1999; Callebaut *et al*. 2002; Dominski *et al*. 2007; Cattell *et al*. 2010). In addition to these domains, Pso2 contains a putative zinc finger motif in its N-terminal region (Cattell *et al*. 2010; this study). The *S. cerevisiae PSO2* encodes a 78 kDa protein, and was initially thought to be specific for ICL repair (Li and Moses, 2003; Cattell *et al*. 2010). However, several lines of evidence suggests that Pso2, and its mammalian counterparts *SNM1A/SNM1B* act in several different pathways and differ in their substrate specificity (Baddock *et al*. 2020). Combined biochemical and structural studies have demonstrated that Pso2/SNM1A/SNM1B exhibit both structure-specific endonuclease and 5’→3’ exonuclease activities; the terminal 5’-phosphate group is essential for the latter activity (Ma *et al*. 2002; Li *et al*. 2005; Hejna *et al*. 2007; Tiefenbach and Junop, 2012; Buzon *et al*. 2018; Baddock *et al*. 2021). Furthermore, a recent study has demonstrated that *S. cerevisiae* Hrq1 helicase stimulates its cognate Pso2 nuclease activity (Rogers *et al*. 2020).

Since clinically relevant concentrations of platinum-based anti-cancer drugs and endogenous aldehydes, under physiologically relevant conditions, cause damage to nuclear and mtDNA by inducing ICLs (Olivero *et al*. 1997; Giurgiovich *et al*. 1997; Cullinane and Bohr, 1998; Brendel *et al*. 2003; Podratz *et al*. 2011), there is substantial interest in exploring whether the enzymes needed for ICL repair exist in the DNA-bearing organelles. Herein, we combined bioinformatics, genetic, microscopic and biochemical approaches with *in vitro* and *in vivo* analyses to assess a previously unrecognized role of targeting signals in the subcellular localization and function of Pso2. Most notably, we found evidence that the nuclear-encoded Pso2 is a dual-localized ICL repair protein, i.e., it can be imported into both mitochondria and the nucleus and that genotoxic agents elevate its abundance in the mitochondria. Through functional analyses, we show that Pso2 contains an N-terminal mitochondrial targeting sequence (MTS) and a pair of nuclear localization signals (NLS1 and NLS2) that are necessary for its import into the mitochondria and nucleus, respectively. Additionally, we noticed that NLS1 is a part of MTS and point mutations within the NLS motifs disrupt the function of Pso2 and its nuclear import; at the same time, they potentiate its enrichment in the mitochondria, indicating an inverse relationship between MTS and NLS motifs. Furthermore, either NLS1 or NLS2 is sufficient for the import of Pso2 into the nucleus; this implies that the two NLS motifs are functionally redundant. Regardless, mutations that attenuate the nucleo-mitochondrial import of Pso2 do not impair its nuclease activity. The potential significance of dual targeting of Pso2 to two DNA-bearing organelles is discussed.

## MATERIALS AND METHODS

### Strains, plasmids and growth conditions

*S. cerevisiae* strains used in this study are described in Table 1. The primer pairs used for PCR amplification are depicted in Table 2. Standard yeast media (YPD, SD), supplemented as noted were used for growth and strain selection (Sherman, 1991). Standard recombinant DNA techniques were used in the construction of recombinant plasmids as described (Sambrook and Russell, 2001). *S. cerevisiae pso2Δ* haploid strain was constructed using a one-step gene replacement method with PCR-generated deletion cassette KanMX4 derived from the plasmid pFA6a-KanMX4 (Janke *et al*. 2004) using primers P1 and P2 (Table 2). The disruption of *PSO2* was confirmed by PCR using primers P3 and P4 (Table 2). *Escherichia coli* strains harbouring plasmids pDEST14-*PSO2* and pDEST14-*pso2*^H611A^ expressing His-tagged wild-type Pso2 and its catalytically-inactive mutant, respectively, were obtained from Murray Junop (Tiefenbach and Junop, 2012). Single (*pso2*_nls1_ and *pso2*_nls2_) and double (*pso2*_nls1-2_) NLS mutants were generated by PCR-based site-directed mutagenesis with pDEST14-*PSO2* as the template. Similarly, the *pso2*_mts_ mutant (pRSETA-*pso2*_mts_) was generated through PCR amplification of the *PSO2 orf* using a pair of mutagenic primers (forward and reverse primers P14 and P6, respectively). The PCR-amplified product (*pso2*_mts_) was inserted into a BamH1/Nco1 site of the pRSET A vector.

**Table 1.**
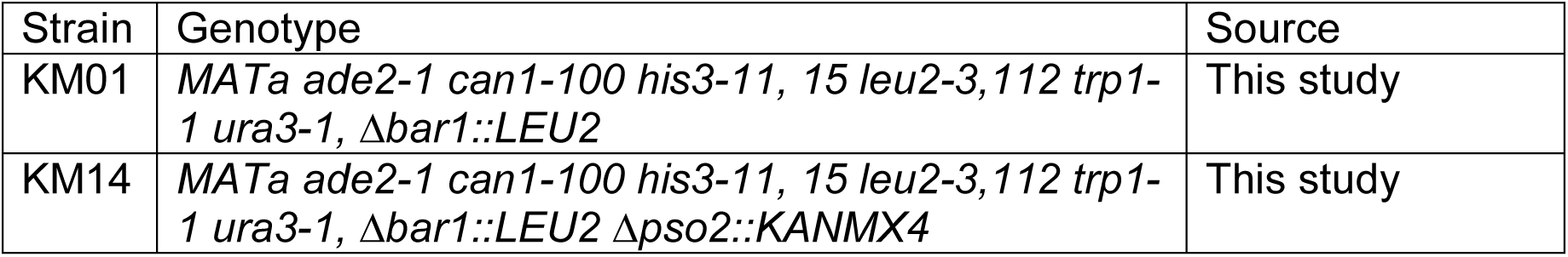
S. cerevisiae strains used in this study

**Table 2.**
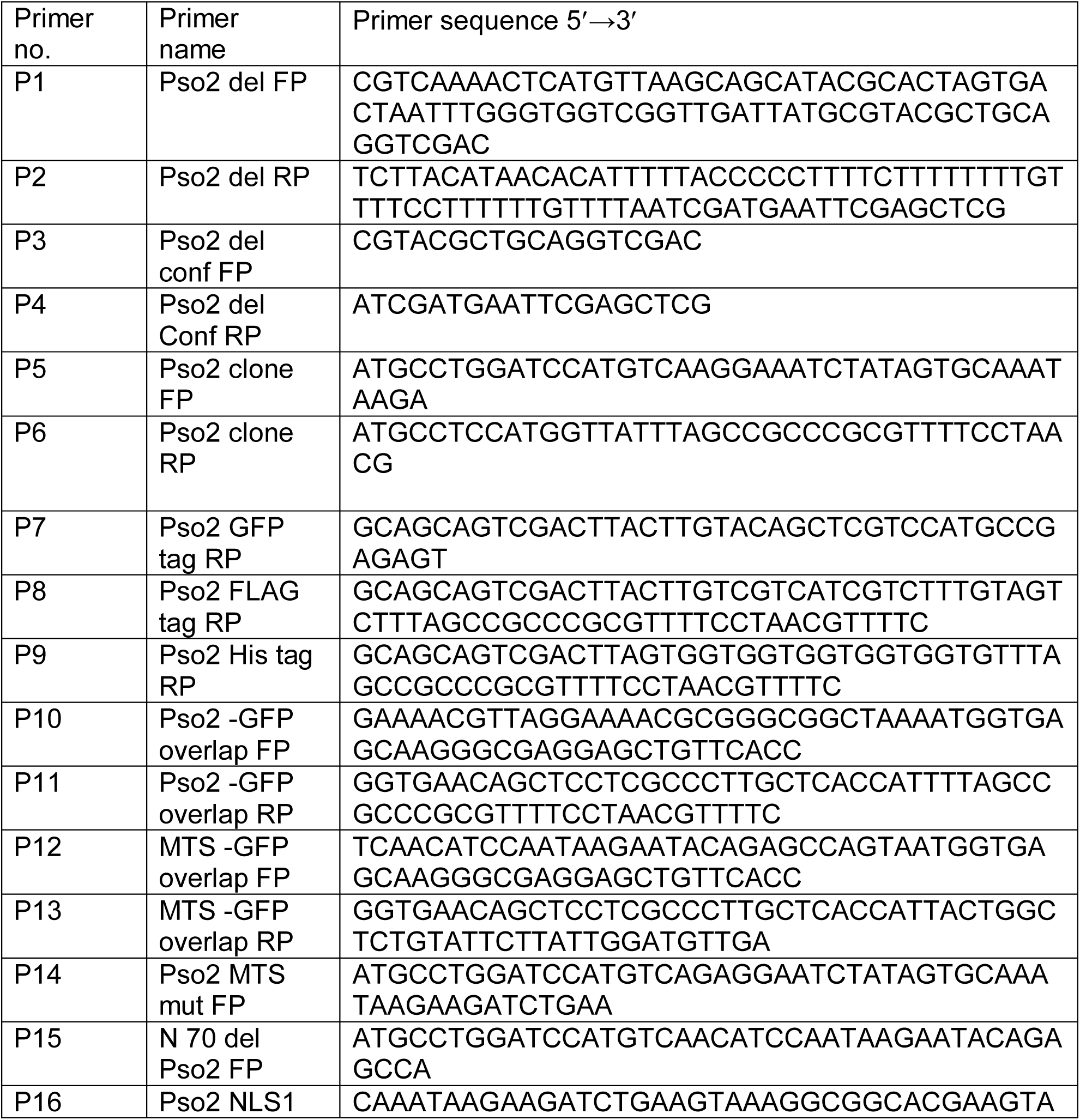
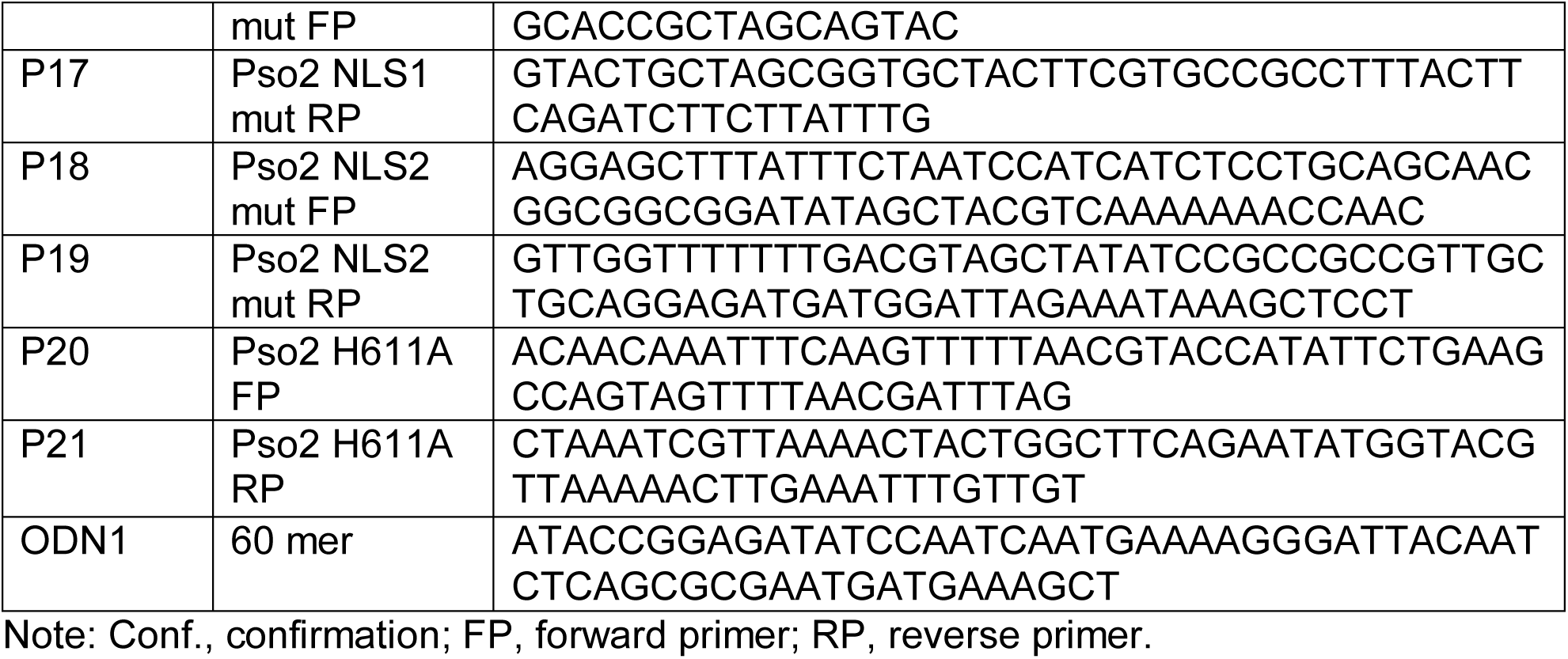
List of primers used for the construction of different pso2 mutants

*S. cerevisiae* strains were grown on YPD plates (1% yeast extract, 2% peptone, 2% dextrose and 2% agar) at 30 °C. Alternatively, the cells were cultured at 30 °C in SD medium containing 0.67% yeast nitrogen base without amino acids, 0.19% yeast synthetic drop-out media supplement containing all essential amino acids minus the amino acid used for selection, 2% dextrose (2% agar for plates). *S. cerevisiae* competent cells were prepared by the LiAc method as described (Knop *et al*. 1999).

### Construction and expression of Pso2-GFP fusion proteins

The Pso2-GFP expression construct was created in two steps using standard molecular cloning procedures. First, the *S. cerevisiae PSO2* ORF was PCR-amplified from the pDEST14-*PSO2* plasmid using primers P5 and P11 (Table 2). In parallel, the gene encoding green fluorescent protein (GFP) in the pcDNA3-EGFP construct was PCR-amplified using primers P10 and P7 (Table 2). In the second step, the two PCR products corresponding to *PSO2* and GFP were fused by overlap extension-PCR. The resulting *PSO2*::*GFP* DNA fragment was inserted into the BamHI-SalI linearized centromeric plasmid pRS416-TEF (Mumberg *et al*. 1995). The resulting expression vector was named as p416-*PSO2*::*GFP*. The C-terminal FLAG-tagged Pso2 construct was generated by PCR amplification of the *PSO2* ORF in the pDEST14-*PSO2* vector using forward primer P5 and reverse primer P8 containing DNA sequence coding for a FLAG tag (Table 2). The resulting *PSO2*::FLAGDNA fragment was inserted into the pRS416-TEF vector. The recombinant construct was designated as p416-*PSO2*::FLAG. This was used as a template to construct the expression vectors p416-*pso2*_nls1_::FLAG, p416-*pso2*_nls2_::FLAG, p416-*pso2*_nls1-2_::FLAG, and p416-*pso2*^H611A^::FLAG by PCR-based site-directed mutagenesis. The p416-*pso2*_mts_::FLAG construct was generated by PCR amplification of *PSO2* insert in the pDEST14-*PSO2* vector using mutagenic primers - P14 and P8 as forward and reverse primers, respectively (Table 2). The resulting *pso2*_mts_ PCR product was inserted into the BamH1/Sal1 site of pRS416-TEF vector.

The p416-*pso2_Δ_*_1-70_::FLAG expression vector was generated by PCR amplification of the *PSO2* ORF using forward primer P15 and reverse primer P8 (Table 2). The PCR-amplified product was inserted into the BamH1/Sal1 site of the pRS416-TEF vector. The plasmid pRS413-MTS::mCherry was obtained from Patrick D’Silva, Indian Institute of Science, Bangalore. This construct harbours DNA sequence coding for MTS from subunit 9 of ATPase synthase fused in-frame with DNA sequence coding for fluorescent protein m-Cherry. The *S. cerevisiae* cells transformed with pRS413-MTS::mCherry plasmid, constitutively express mitochondria-targeted mCherry (Bankapalli *et al*. 2015).

### Confocal laser scanning microscopy

The *S. cerevisiae pso2Δ* cells were co-transformed with recombinant p416-*PSO2*::GFP and p413-MTS::mCherry plasmids. The transformants were selected on synthetic medium supplement without uracil and histidine. The transformant coexpressing Pso2-GFP and MTS-mCherry were grown in 20 ml of SD medium lacking uracil and histidine at 30 °C until the exponential phase. The cells in the culture were evenly distributed into four 50 ml Erlenmeyer flasks: first one served as an untreated control and others were incubated with either 500 µM cisplatin, 0.1% MMS or 5 mM H_2_O_2_ for 2 h at 30 °C. The cells were harvested by centrifugation and the pellet was washed with 1x PBS. The pellet was resuspended in 1x PBS and cells were stained with a solution containing 1 µg/ml 4,6-diamidino-2-phenylindole (DAPI) (Sigma-Aldrich India) for 10 min. A Fluoview FV 3000 Confocal Laser Scanning microscope (Olympus) was used to visualize the cells. The images were captured and processed using the CellSens Software (DSS Imagetech Private Limited, Delhi).

### Isolation of *S. cerevisiae* mitochondria

The mitochondria were isolated from *S. cerevisiae* whole cell lysates as described (Meisinger *et al*. 2006). Briefly, exponentially growing cells (1 L) were harvested by centrifugation at 3000 x g for 5 min at 24 °C. The cell pellet was resuspended in buffer (100 mM Tris-H_2_SO_4_ buffer, pH 9.4, containing 10 mM DTT) and incubated at 30 °C for 20 min with gentle shaking (80 rpm). Cells were collected by centrifugation at 3000 x g for 5 min and resuspended in a buffer (20 mM potassium phosphate buffer, pH 7.4, with1.2 M sorbitol) containing 3 mg zymolyase (US Biologicals) per gram wet weight of the pellet. The cell suspension was incubated at 30 °C for 45 min with gentle shaking to enhance spheroplast formation. The cells treated with zymolyase were centrifuged at 3000 x g for 5 min at 4 °C. The pellet was resuspended in cold homogenization buffer (10 mM Tris-HCl buffer, pH 7.4, 0.6 M sorbitol, 1 mM EDTA, 1 mM phenylmethylsulfonyl fluoride and 0.3% (w/v) bovine serum albumin). The spheroplasts were lysed using the Dounce homogenizer and the homogenate was centrifuged at 1500 x g for 5 min to remove cell debris and nuclei. The supernatant was centrifuged at 4000 x g for 5 min. The supernatant from the previous step was centrifuged at 12,0 00 x g for 20 min. The pellet was resuspended in cold SEM buffer (10 mM MOPS-KOH, pH 7.2, buffer with 250 mM sucrose and 1 mM EDTA). The suspension of mitochondria was placed on the top of a discontinuous sucrose gradient (1.5 ml 60%, 4 ml 32%, 1.5 ml 23%, and 1.5 ml 15% sucrose) in EM buffer (10 mM MOPS-KOH buffer, pH 7.2, containing 1 mM EDTA). Centrifugation was performed in a SW 28 swinging-bucket rotor (Beckman) at 134,000 g for 1 h at 2 °C. The intact mitochondria formed a brown band at the 60% and 32% sucrose interface. The purity of mitochondrial preparation was confirmed by the absence of ER/PM marker. The isolated mitochondria were resuspended in SEM buffer and stored at -80 °C.

### Isolation of *S. cerevisiae* nuclei

*S. cerevisiae* nuclei were isolated from whole cell lysates as described (Sperling and Grunstein, 2009). Briefly, cells (2 L) at an optical density (OD_600_) of 2 were harvested by centrifugation at 5000 rpm for 5 min at 24 °C. The pellet was washed once with 1x PBS and re-suspended in 35 ml of zymolyase buffer (50 mM Tris-HCl buffer, pH 7.5, containing 1 M sorbitol, 10 mM MgCl_2_ and 3 mM DTT). The cells were incubated at 30 °C for 15 min with gentle shaking. Cells were collected by centrifugation and the pellet was resuspended in 30 ml zymolyase buffer to which 20 ml of YPD/S (YPD medium containing 1 M sorbitol) was added. The samples were incubated with 15 mg zymolyase at 30°C with gentle shaking to facilitate spheroplast formation. After 90 min incubation, 100 ml YPD/S medium was added to the samples and the cells were collected by centrifugation at 4 °C. The pellet was washed once with 250 ml cold YPD/S medium and then with 200 ml cold 1 M sorbitol, then resuspended in 100 ml ice cold buffer N (30 mM HEPES buffer, pH 7.6, containing 25 mM K_2_SO_4_, 5 mM MgSO_4_, 1 mM EDTA, 10% glycerol, 0.5% Nonidet P-40, 7.2 mM spermidine, 3 mM DTT, and 1x protease inhibitor cocktail). The spheroplasts were lysed using the Yamamoto homogenizer set at 100 rpm. The lysate was centrifuged at 2000 rpm for 10 min at 4 °C in a JA-20 rotor (Beckman). The supernatant from the previous step was centrifuged in a Beckman Coulter high speed centrifuge equipped with a JA-20 rotor at 6000 rpm for 20 min at 4 °C. The pellet was washed once with buffer N and resuspended in 1 ml of buffer N. Protein concentration was determined using a Nano drop spectrophotometer by diluting 10 µl sample in 990 µl of 2 M NaCl. The aliquots were stored at -80 °C.

### Protease protection assay

The assay was carried out as described (Liu *et al*. 2015). Briefly, aliquots containing isolated mitochondria (20 µg) in SEM buffer (10 mM MOPS-KOH buffer, pH 7.2, containing 250 mM sucrose and 1 mM EDTA) were placed in four 1.5 ml Eppendorf tubes. While the first aliquot served as a control, the second was treated with proteinase K (10 µg*/*ml), the third with 1% Triton X-100 and the fourth with 10 µg*/*ml proteinase K and 1% Triton X-100. After incubation on ice for 30 min, the reaction was stopped by adding phenylmethyl sulfonyl fluoride (Sigma Aldrich India) to a final concentration of 5 mM. The samples were centrifuged at 12000 x g for 30 min at 4 °C to collect mitochondria. The pellet was resuspended in SDS-polyacrylamide gel electrophoresis (SDS-PAGE) sample buffer, followed by boiling at 95 °C for 10 min. The samples were separated on a 10% SDS-PAGE and subjected to Western blot analysis.

### Alkali extraction of mitochondria

The assay was performed as described (Bannwarth *et al*. 2012). Briefly, aliquots containing isolated mitochondria (20 µg) were added to 100 µl SEM buffer (10 mM MOPS-KOH buffer, pH 7.2, 250 mM sucrose,1 mM EDTA and 100 mM Na_2_CO_3_, pH 11.5). After incubation at 4 °C for 30 min, the samples were centrifuged at 14000 x g for 30 min at 4 °C. The supernatant was transferred to a fresh tube. The pellet was washed once with SEM buffer and resuspended in 200 µl of SEM buffer. Aliquots from both supernatant and pellet fractions were separated on a 10 % SDS-PAGE and subjected to Western blot analysis.

### Preparation of mitoplasts

An aliquot of isolated mitochondria (20 µg) was diluted 10-fold with hypotonic buffer (10 mM MOPS-KOH buffer, pH 7.2). After incubation for 20 min at 4 °C, the samples were centrifuged at 14000 x g for 30 min at 4 °C. The pellet was resuspended in SEM buffer (10 mM MOPS-KOH buffer, pH 7.2, containing 250 mM sucrose and 1 mM EDTA). Aliquots were separated on a 10% SDS-PAGE and subjected to Western blot analysis.

### Western blot analysis

The protein samples (15–20 μg) from whole cell lysates, nuclear and mitochondrial fraction were separated by 10-12% SDS-PAGE and transferred onto PVDF membranes as described (Mahmood and Yang, 2012). The membranes were blocked for 1 h at 24 °C or 6 h at 4 °C with 5% non-fat dry milk in TBST buffer (20 mM Tris-HCl buffer, pH 7.5, containing 150 mM NaCl and 0.1% Tween 20).The membranes were probed with the indicated primary antibody in TBST buffer containing 3% bovine serum albumin. The antibodies were used in the following dilutions: anti-FLAG (1:5000), anti-Tim23 (1:5000), anti-Tom70 (1:5000), anti-Mge1(1:6000), anti-Ydj1(1:6000), anti-H3 (1:4000), and anti-Pgk1 (1:2000) at 4 °C for 12 h. The membranes were washed 3 times for 10 min each with TBST buffer following incubation with either horseradish peroxidase-conjugated secondary anti-rabbit (1:1000) or anti mouse (1:8000) antibody for 1 h at 24 °C. Finally, the membranes were washed 3 times for 10 min each with TBST buffer at 24 °C and visualized using enhanced chemiluminescence reagents (Bio-Rad Laboratories, CA, USA). The images were captured using Image Quant LAS 4000 (GE Healthcare Life Sciences, PA, USA). The polyclonal antibodies against Tim23, Tom70, Mge1, and Ydj1 were provided by Patrick D’Silva of the Indian Institute of Science, Bangalore; the antibodies against other proteins were obtained from Sigma-Aldrich India, except anti-Pgk1, which was from Santa Cruz Biotechnology, CA, USA. The signals detected by Western blot analysis were quantified using imageJ software.

### Complementation assay

*S. cerevisiae pso2Δ* cells harbouring either empty vector, recombinant plasmids expressing the wild-type Pso2 or its mutant variants were grown until the exponential phase in 10 ml in uracil-lacking SD medium. Each culture was divided into two equal halves: one served as an untreated control and the second was treated with cisplatin at a final concentration of 300 µM. After incubation for 2 h at 30 °C, the cells were harvested by centrifugation and the pellet was washed once with 1x PBS. The pellets were resuspended in 1x PBS and the optical density of cell suspension was normalized to an OD_600_ of 1.0. Five µL aliquots were spotted from the serial dilutions on uracil-lacking SD medium plates. The plates were incubated at 30 °C, and pictures were taken after 72 h.

### Overexpression and purification of the wild-type and variant species of Pso2

The recombinant Pso2 was purified as described (Tiefenbach and Junop, 2012). Briefly, the *E. coli* BL21 (DE3) expression host strain (Novagen) was transformed with the pDEST14-*PSO2* expression vector. Cells grown in LB broth to an OD_600_ of 0.5 at 37 °C, the Pso2 expression was induced with the addition of 1mM isopropyl β-D-thiogalactopyranoside. After incubation at 16 °C for 16 h with gentle stirring, the cells were harvested by centrifugation at 3000 x g for 10 min at 4 °C. The pellet was washed once with buffer (20 mM sodium phosphate buffer, pH 7.0, containing 200 mM NaCl and 10% (v/v) glycerol) and resuspended in buffer B (50 mM sodium phosphate buffer, pH 7.0, containing 500 mM NaCl, 3 mM β-mercaptoethanol and 10% (v/v) glycerol). Cells were lysed using an ultrasonic processor set to 60% amplitude for 5 min and centrifuged at 72000 x *g* for 90 min. The supernatant was applied to a Ni^2+^-NTA affinity column (5 ml bed vol) pre-equilibrated with buffer C (50 mM sodium phosphate buffer, pH 7.0, containing 500 mM NaCl, and 10% (v/v) glycerol). The column was washed with buffer C containing 45 mM imidazole. The bound protein was eluted with a linear gradient of 45 → 250 mM imidazole in buffer C. The fractions containing Pso2 were pooled and dialyzed against buffer D (50 mM sodium phosphate buffer, pH 7.0, containing 100 mM NaCl, 5 mM DTT and 10% (v/v) glycerol). The dialysate was applied to a Q-Sepharose column which had been equilibrated with buffer D. The bound protein was eluted with a linear gradient of 100 → 200 mM NaCl in buffer D. The fractions containing Pso2 were combined and dialysed against a storage buffer (10 mM Tris-HCl buffer, pH 7.5, containing 100 mM NaCl, 5 mM DTT and 10% glycerol). During all steps of purification, dye-binding assay was used for the determination of protein concentrations (Bradford, 1976) and the purity was assessed by 12% SDS-PAGE (Green and Sambrook, 2012). The total yields of purified wild-type Pso2 and its variants was approximately 2.1 mg from 5 L culture. Aliquots were stored at -80 °C. The Pso2_nls_ and Pso2_mts_ mutants were purified using the same protocol as that of wild-type Pso2 and stored at -80 °C.

### Nuclease assay

The assay was performed as described (Tiefenbach and Junop, 2012). Briefly, a 60-mer oligonucleotide (Table2) was labelled at the 5’ end with [γ-^32^P]ATP and T4 polynucleotide kinase as described (Green and Sambrook, 2012). The reaction mixture (20 µl) containing 10 mM Tris-HCl, pH 7.9, 10 mM MgCl_2_, 50 mM NaCl, and 1 mM DTT and 3 nM ^32^P-labeled 60-mer single-stranded DNA (ssDNA) was incubated with varying concentrations of the wild-type Pso2 or its mutant at 37 °C for 30 min. The reaction was stopped by adding gel loading dye (95% formamide, 5 mM EDTA, 0.025% bromophenol blue and 0.025% xylene cyanol); the sample were incubated at 95 °C for 10 min and, subsequently, at 4 °C for 5 min. The samples were separated on a 15% denaturing gel by electrophoresis in 89 mM Tris-borate buffer, pH 8.3. The gels were dried and exposed to a phosphor imaging screen. The images were acquired using the Fuji FLA-5000 phosphor imaging system.

### Bioinformatics analysis

The mitochondrial targeting signals in Pso2 were identified using the prediction algorithm MitoProt II, and PSORT II. Similarly, the cNLS mapper, and PSORT II prediction algorithm were used to identify nuclear localization signals in Pso2.

### Statistical analyses

Statistical analyses were carried out using GraphPad Prism 6 software. The statistical parameters are expressed, and the corresponding p-values and sample size are described in the Fig. legends. Statistical analysis between two groups was done by paired or unpaired two-tailed t-test.

## RESULTS

### *S. cerevisiae* Pso2 contains an N-terminal MTS and two NLS motifs

An earlier study demonstrated localization of Pso2 to the cell nucleus through a high-throughput microscopy screen of the *S. cerevisiae* GFP fusion collection (Tkach *et al*. 2012); however, it remained unknown whether its import is autonomous or mediated by nuclear localization signal(s). Furthermore, the roles of cell-intrinsic (e.g., genetic and physiological pathways) and cell-extrinsic (e.g., nutrient signals, genotoxic and mechanical stresses) factors that abet or impede the subcellular localization of Pso2 have not been identified to date. Additionally, how the subcellular localization of Pso2 might contribute to its physiological functions has not been addressed. In the absence of such knowledge, we used bioinformatics tools to identify the presence of subcellular protein targeting signals in Pso2. Signal sequence analysis of the Pso2 amino acid sequence using PSORT II algorithm yielded strong score of 0.84 for NLS. A similar analysis using MitoProt II algorithm yielded a score of 0.88 for MTS. This implies that Pso2 contains putative MTS and NLS motifs. The *S. cerevisiae* Pso2 is a 661 amino acid polypeptide, which can be divided schematically into two regions for the purpose of the current study; the N- and C-terminal regions (Fig. 1A). A bioinformatics analysis of the Pso2 primary sequence for conserved motifs flagged the existence of one MTS, a pair of NLSs (NLS1 and NLS2) and a zinc-finger domain all of them are located between 1-200 amino acid residues at its N-terminus. Within this region, while amino acid residues 1-70 correspond to MTS, residues 15-18 and 194-200 to the NLS1 and NLS2 motifs, respectively. A similar scan flagged that NLS1 is located within MTS at its proximal end, thereby suggesting a potential interplay between these two motifs.

**Fig. 1.**
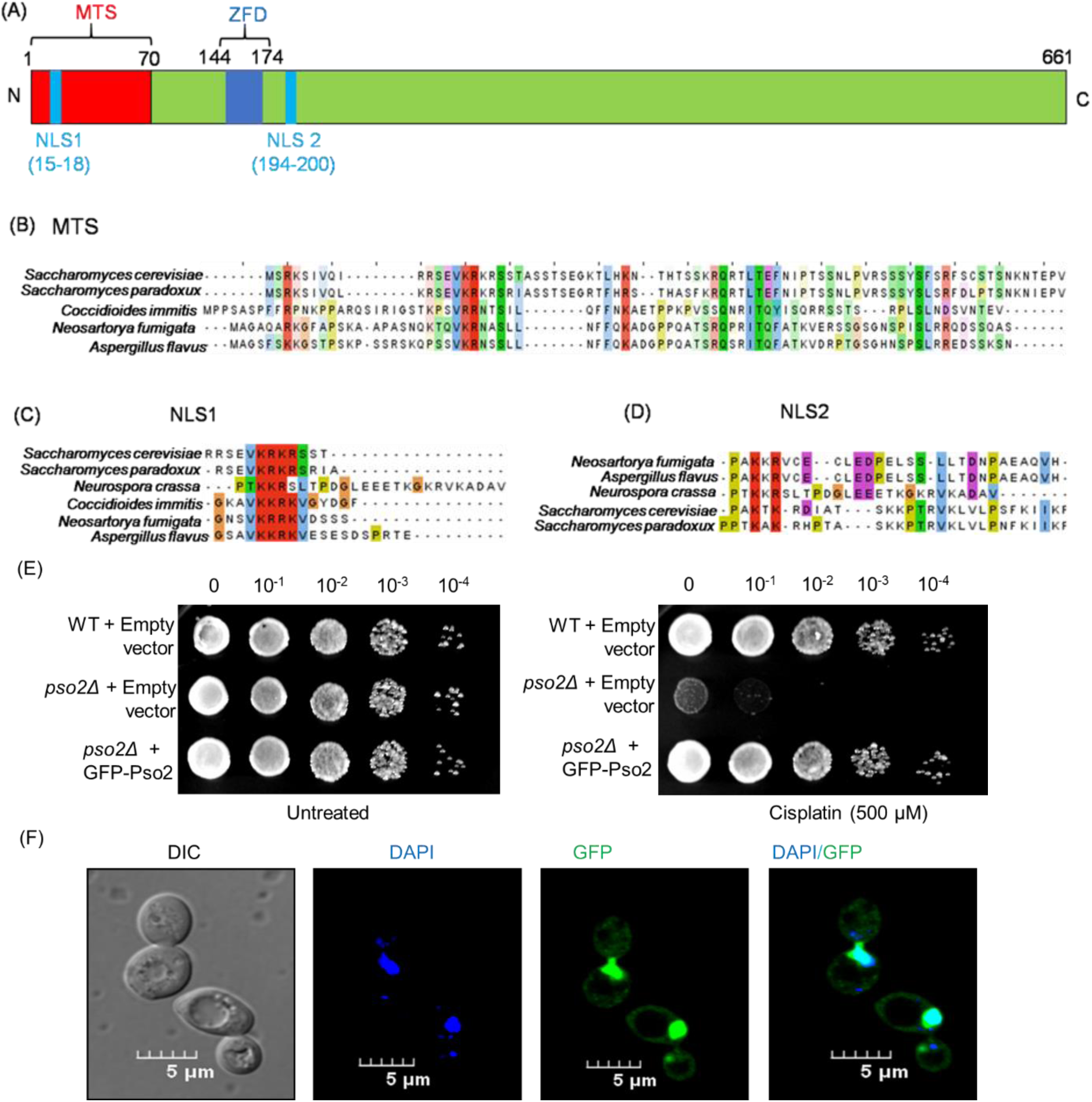
*S. cerevisiae* Pso2 harbours a MTS and two NLS sequence motifs, and protects the *pso2Δ* strain from cisplatin-induced cell death. (A) Domain organization of Pso2 showing putative MTS, NLS1, NLS2 and a zinc-finger domain (ZFD). Numbers represent the positions of the amino acid residues in the indicated motifs. N and C represent the N- and C-terminal ends, respectively. (B) A multiple sequence alignment of MTS sequence in five fungal species. (C) and (D) Multiple sequence alignments of NLS1 and NLS2 sequences, respectively. The highly conserved residues are highlighted in red in the sequence alignment. The MTS and NLS sequences were detected using MitoProt II, and cNLS Mapper tools, respectively. The conserved sequences were analysed for conservation by Clustal Omega and then viewed by Jalview. ClustalX colour code is as follows, blue for hydrophobic, green for polar, magenta and red for negative and positively charged residues, respectively; cyan for aromatic, yellow for prolines and orange for glycines. Amino acid sequence of *S. cerevisiae* Pso2 was retrieved from *Saccharomyces* Genome Database (SGD), and amino acid sequences of Pso2 from *S. paradoxus*, *Coccidioidesimmitis, Neosartoryafumigata, N.crassa* and *A. flavus* were retrieved from UniProt. (E) Complementation of *S. cerevisiae pso2Δ* cells with Pso2::GFP completely attenuates cisplatin-induced cell death. The *S. cerevisiaepso2Δ*strain was transformed either with empty vector or p416-*PSO2*::GFP construct. The assay was performed as described in Methods. (F) Subcellular localization of Pso2-GFP in *S. cerevisiae pso2Δ* cells. The CLSM images, from left to right: DIC image, DAPI image, Pso2-GFP image, and DAPI-GFP merged image. The images in panel B and C are representative of three independent experiments.

### MTS and NLS sequences are conserved among fungal species

Typically, the NLS sequences contain one (monopartite) or two (bipartite) clusters of positively charged amino acid residues, each bearing two to four basic amino acid moieties (either Lys or Arg). As mentioned above, Pso2 possesses an N-terminal MTS and two NLS motifs; NLS1 and NLS2 (Fig. 1A). A multiple sequence alignment of the deduced amino acid sequence of the MTS domain from five different fungal species enabled the identification of well-conserved repetitive clusters of positively charged amino acid residues across the 70-amino acid region (Fig. 1B). Likewise, alignment of the amino-acid sequence of Pso2 orthologs revealed that the canonical bipartite NLS1 motif is highly conserved in the Pso2 orthologs of six fungal species examined in this study (Fig. 1C). On the contrary, NLS2 motif shows a substantially lower level of sequence conservation (Fig. 1D). These results are consistent with different protein families that possess non-canonical NLS motifs, including the recently discovered class of functional bipartite NLS motifs (Koch et al, 2018; Makkerh et al, 1996). Thus, the conserved features of Pso2 MTS and NLS motifs prompted us to investigate their ability to function as organelle targeting signals *in vivo*. In addition, we asked whether either one or both NLSs are essential for targeting Pso2 into the nucleus.

### Functional validation of Pso2-GFP fusion construct

We first sought to determine the function and subcellular localization of Pso2 in cells grown under normal conditions. To investigate this, (as Pso2 is a low abundance protein) a plasmid (p416-*PSO2*::GFP) that allows constitutive expression of Pso2::GFP was constructed; this makes it possible to visualize its localization using confocal laser scanning microscopy (CLSM). A complementation assay was performed to ascertain the *in vivo* functional activity of GFP-tagged Pso2 in the *pso2Δ* mutant background. In agreement with our rationale, while GFP-tagged Pso2 could rescue the cisplatin hypersensitive phenotype of *pso2Δ* cells, deletion of *PSO2* exacerbated cisplatin-induced cell death (Fig. 1E). To correlate these results, its subcellular localization was visualized using a CLSM. In accordance with a previous report (Tkach et al. 2012). We found localization of GFP-tagged Pso2 to the DAPI-stained nuclei of unstressed cells and, more importantly, Pso2-GFP puncta were also found throughout the cytoplasm (Fig. 1F). Altogether, these results suggest that the GFP-tagged Pso2 behaves similar to that of the wild-type protein *in vivo*.

### *S. cerevisiae* Pso2 localizes to the mitochondria of unstressed cells

Motivated by the observation that a significant fraction of Pso2-GFP puncta were found in the cytoplasm (Fig. 1F), we carefully examined its localization to mitochondria in unstressed cells. To do this, a plasmid expressing Pso2-GFP was co-transformed with MTS-mCherry expression construct into *S. cerevisiae pso2Δ* cells, and the cells were visualized using CLSM. As anticipated, the vast majority of Pso2-GFP was found in the nucleus (Fig. 2A). Importantly, the same cells showed colocalization of Pso2-GFP and MTS-mCherry fluorescent foci, indicating the presence of Pso2 in the mitochondria (Fig. 2A). As such, strong colocalization of Pso2-GFP puncta with MTS-mCherry could not be attributed to fluorescence noise or spread caused by fixation or imaging artefacts since no obvious diffusion of DAPI or mCherry fluorescence was seen in these cells. To validate this notion, a biochemical approach was utilized to assess the presence of Pso2 in the mitochondria. A Western blot analysis of whole-cell lysates from the isolated mitochondria indeed confirmed the presence of Pso2 in the mitochondria (Fig. 2B, 2C). More importantly, this analysis demonstrated the presence of both 78 kDa and 68 kDa species of Pso2 (Fig. 2B, lane 1). In parallel, similar analysis of lysates using antibodies against Ydj1 and histone H3 informed the purity of the mitochondrial fraction and lack of cross-contamination from cytosolic and nuclear fraction, respectively.

**Fig. 2.**
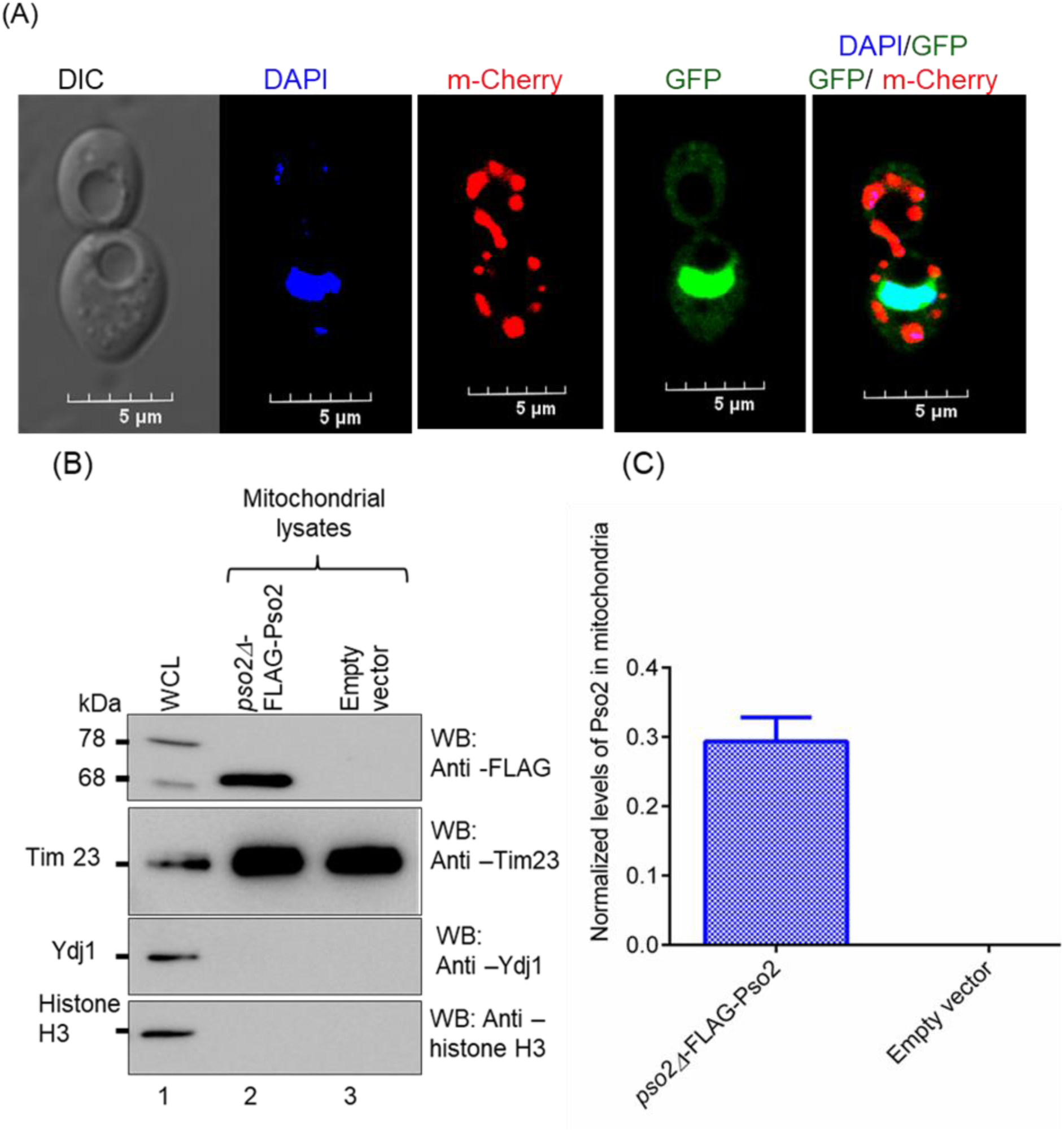
Pso2 localizes to the mitochondria under normal growth conditions. (A) Subcellular localization of Pso2-GFP in *S. cerevisiae pso2Δ* cells. The panels from left to right depict DIC, DAPI, mCherry, GFP, merged DAPI/GFP and GFP/mCherry confocal images, respectively. (B) Western blot analysis mitochondrial lysates of *S. cerevisiae pso2Δ* cells harbouring the plasmid expressing Flag-tagged Pso2 or empty vector. Lane 1, whole cell lysate; 2, mitochondrial lysate; 3, empty vector. The three lower panels were probed with anti-Tim23, anti-Ydj and anti-histone H3 antibodies, respectively. A total of 15 μg protein was loaded per lane for each sample. (C) Representative histograms showing the Pso2 levels in the mitochondria. The error bar represents mean ± SEM from three independent experiments.

### Stress-inducing agents markedly enhance Pso2 levels in the mitochondria

As outlined in the Introduction, several lines of evidence indicate that stress-inducing agents inflict damage to mitochondrial DNA (Contamine and Picard, 2000; Acevedo-Torres *et al*. 2009; Shokolenko *et al*. 2009; Marullo *et al*. 2013; Yimit *et al*. 2019). Given the fact that Pso2-GFP was distributed uniformly throughout the nucleus, and weaker Pso2-GFP puncta were seen in the cytoplasm of unstressed cells (Fig. 1F), we speculated that Pso2 import into the mitochondria might increase following treatment with stress-inducing agents. We tested this hypothesis by co-transforming the *S. cerevisiae pso2Δ* strain with plasmids expressing GFP-tagged Pso2 and MTS-mCherry reporter; the latter is a demonstrated robust mitochondrial matrix marker (Bankapalli *et al*. 2015).

The resulting *pso2Δ* strain expressing GFP-tagged Pso2 and MTS-mCherry reporter was grown to mid-exponential phase and then exposed to diverse genotoxic agents, including methyl methanesulfonate (MMS), cisplatin and H_2_O_2_ at 30 °C. Two hours after drug addition, the cells were imaged using a CLSM. As expected, red fluorescent protein mCherry predominantly localized to the mitochondria in the perinuclear region, which was clearly distinguishable from nuclei stained with DAPI (Fig. 3 A-C, compare column 3 with column 2, from left to right). Likewise, Pso2-GFP fluorescent foci were also seen around the perinuclear region (Fig. 3 A-C, column 4). Importantly, the merged images revealed evidence for extensive colocalization of the red fluorescence of MTS-mCherry with GFP-Pso2, and the fluorescent signal of DAPI with GFP-Pso2 in cells subjected to genotoxic stress (Fig. 3A-C, column 5). Moreover, such colocalization was more robust in MMS-treated cells, as compared with cells exposed to either cisplatin or H_2_O_2_. In line with this, quantitative analysis of CLSM data followed by statistical analysis of colocalization (Manders et al, 1993) showed a 3.5- fold increase in the levels of Pso2 in the mitochondria of cells exposed to MMS, while 2.5- and 2-fold increase was observed in cells treated with cisplatin and H_2_O_2_, respectively, relative to untreated cells (Fig. 3D).

**Fig. 3.**
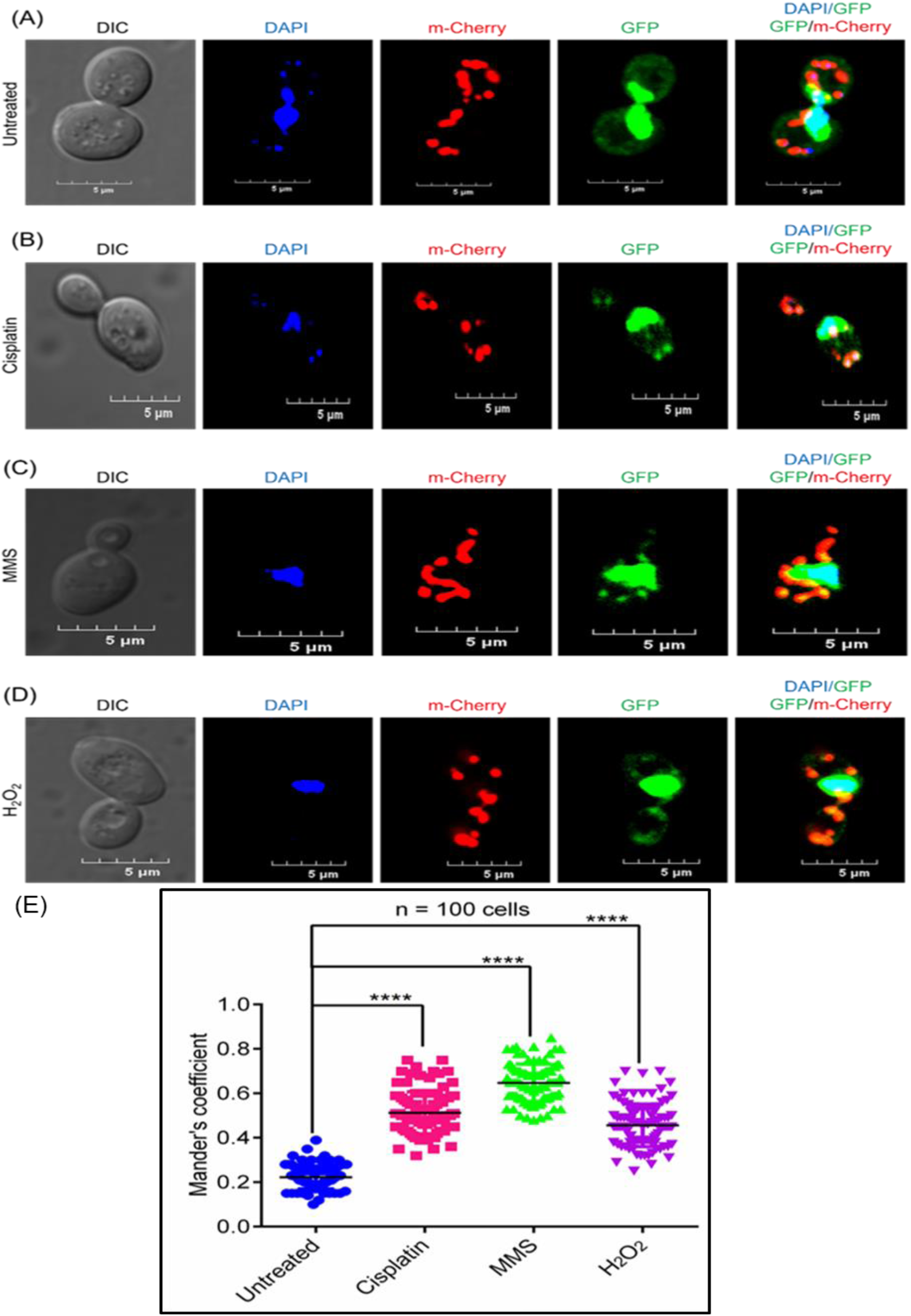
Genotoxic agents enhance the mitochondrial localization of Pso2. Representative CLSM images of *S. cerevisiae pso2Δ* cells expressing Pso2-GFP and MTS-mCherry. (A) Untreated cells. After treatment with (B) 500 µM cisplatin; (C) 0.1% MMS or (D) 5 mM H_2_O_2_ for 2 h at 30 °C. (E) Scatter plot depicting the colocalization of Pso2-GFP foci with MTS-mCherry, calculated using JaCop in Image J software. n=100 cells refers to the total number of cells used for calculating the Manders’ colocalization coefficient values. The mean of three experiments is indicated by a horizontal black line. Statistical analysis was done by un paired Student’s t test (ns, not significant, ****, P < 0.0001).

As further validation that genotoxic stress induces targeting of Pso2 to the mitochondria, Western blot analysis was performed to determine the levels of Pso2 in the mitochondria and nucleus after treatment with various genotoxic agents. Consistent with CLSM data, Western blot analysis of lysates from purified mitochondria and nucleus demonstrated a 2- to 3-fold increase in the levels of Pso2 in the mitochondria of cells treated with genotoxic agents, as compared to untreated cells (Fig. 4A-B). On the contrary, statistically, no significant changes in the levels of Pso2 was observed in the nuclear lysates (Fig. 4C-D). Interestingly, immunoblotting results for the whole cell lysates revealed a faster-migrating protein band of 68 kDa, in addition to the expected 78 kDa species of C-terminally Flag tagged-Pso2 (Fig. 34, lane 1). This result is consistent with the idea that MTS at the N-terminus of Pso2 is cleaved off after import into the mitochondria, thereby yielding a truncated species of 68 kDa. Tim23, Ydj1 and histone H3 were used as mitochondrial, cytosolic and nuclear markers, respectively, to ascertain contamination during cell fraction isolation. To summarize, confocal microscopy combined with quantitative and statistical colocalization analysis, and Western blot data support the view that genotoxic agents differentially modulate the import of Pso2 into the mitochondria, while nuclear import was not affected.

**Fig. 4.**
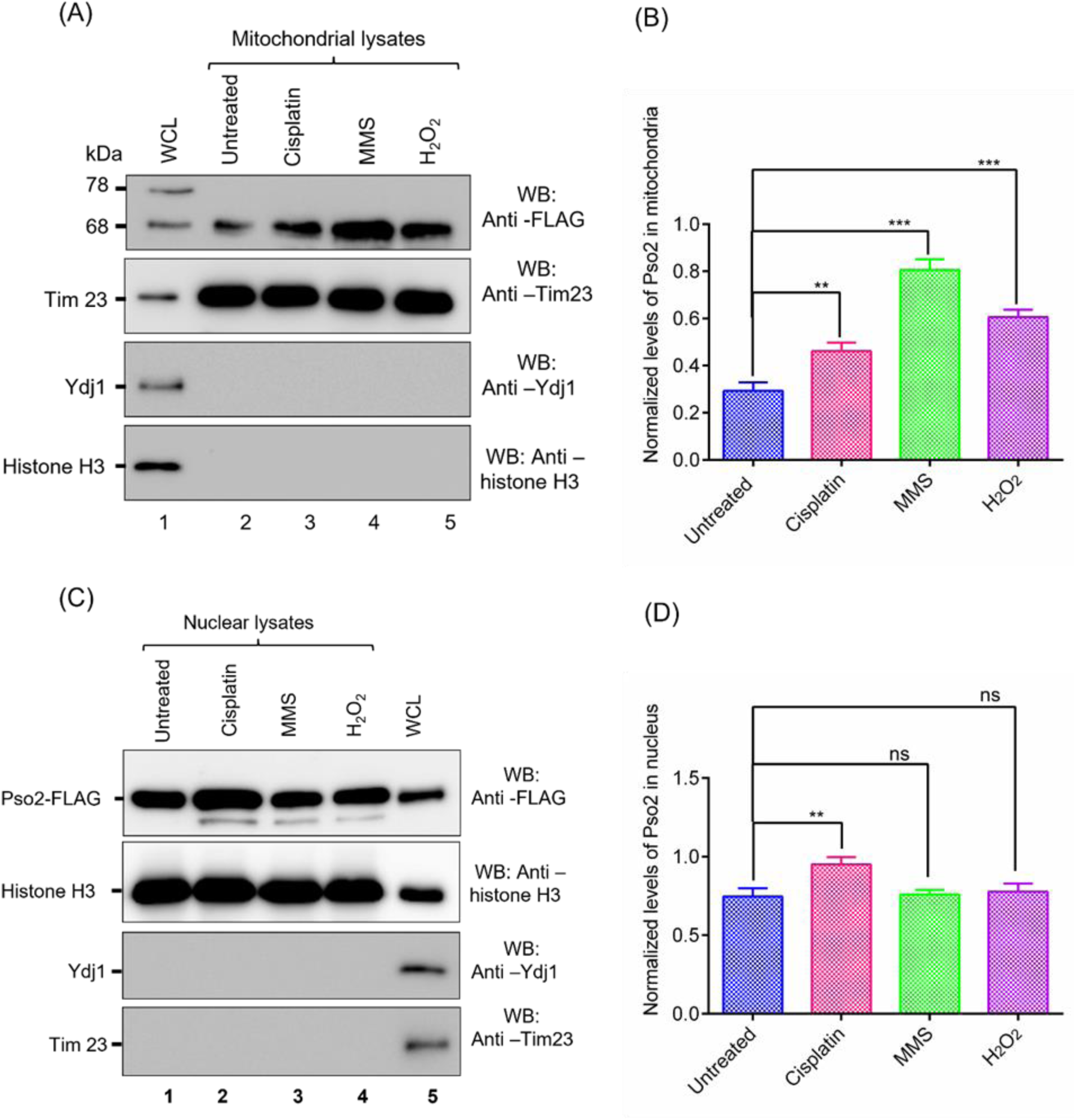
Genotoxic agents differentially modulate mitochondrial localization of Pso2, with no effect on nuclear import. (A) Western blot analysis of mitochondrial lysates. Lane 1, whole cell lysate from untreated cells; 2. mitochondrial lysate from untreated cells. Lanes 3-5, mitochondrial lysates of cells exposed to cisplatin (500 µM), MMS (0.1%) and H_2_O_2_ (5 mM), respectively. A total of 15 μg protein was loaded per lane for each sample. The three lower panels were probed with anti-Tim 23, anti-Ydj and anti-histone H3 antibodies, respectively. (B) Histograms show genotoxin-induced Pso2 levels in the mitochondria, normalized to Tim23 loading control. (C) Western blot analysis of nuclear lysates. Lane 1, nuclear lysate from untreated cells. Lanes 2-5, nuclear lysates of cells exposed to cisplatin (500 µM), MMS (0.1%) and H_2_O_2_ (5 mM), respectively. Lane 5, whole cell lysate from untreated cells. A total of 15 μg protein was loaded per lane for each sample. The three lower panels were probed with anti-histone, anti-Ydj and anti-Tim23 antibodies, respectively. (D) Histograms show genotoxin-induced Pso2 levels in the nucleus normalized to histone H3 loading control. The quantitative data in (B) and (D) was obtained from quantification of band intensities of three independent Western blots. Statistical comparisons were performed by using unpaired Student’s t test (*p < 0.05, **p < 0.01, ***p < 0.001, ns, not significant)

### Either NLS1 or NLS2 is sufficient for nuclear import of Pso2

A plethora of studies reveal that basic amino acid residues play a pivotal role in targeting proteins to the nucleus or mitochondria (Martin *et al*. 2015). To systematically validate the bioinformatics data, mutations were introduced in the putative NLS motifs of Pso2. The conserved amino acid residues in the NLSs are: Arg^16^ and Lys^17^ in the NLS1 and Lys^196^, Lys^198^ and Arg^199^ in the NLS2 motifs respectively (Table 3). The mutant constructs were generated using the p416-*PSO2::FLAG* plasmid as the template by substituting these residues with alanine in the NLS1, NLS2 motifs or both through PCR-based site-directed mutagenesis. In parallel, we also constructed a plasmid bearing point mutations of the conserved residues (Arg3Ala and Lys4Ala) in the MTS motif (Table 3).

**Table 3.**
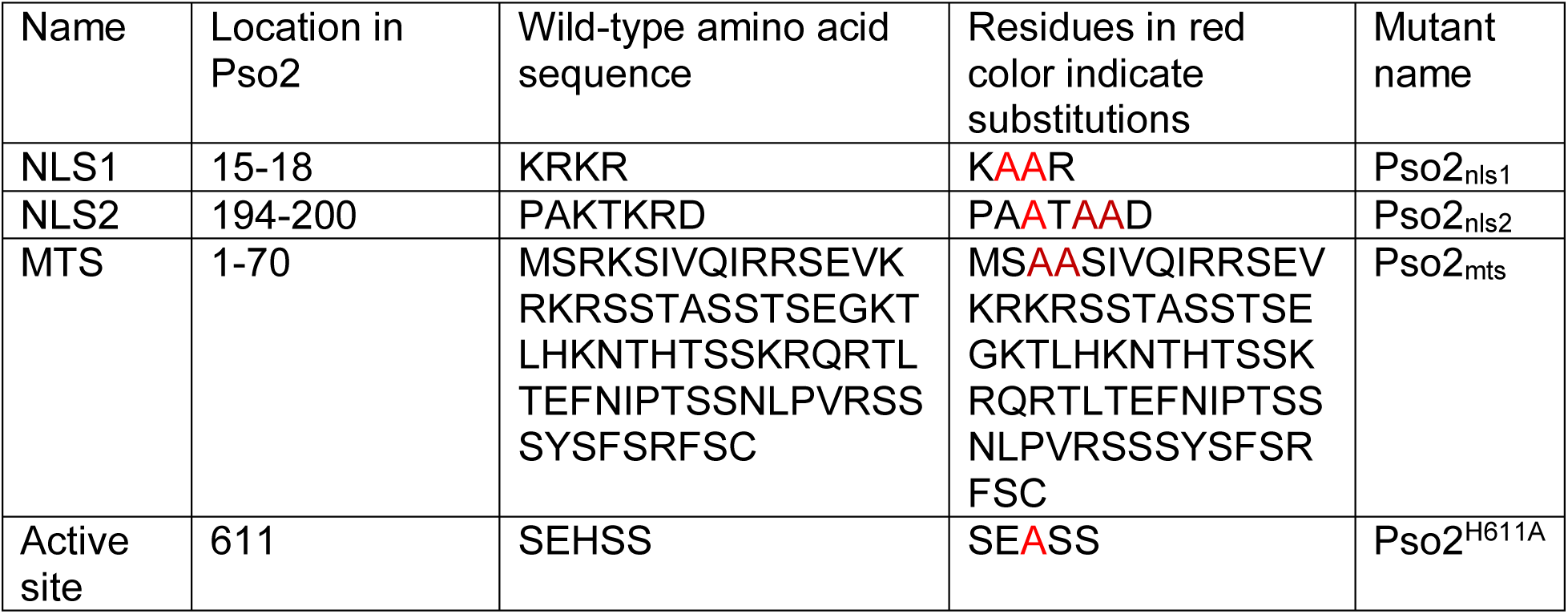
NLS1, NLS2, MTS and active site amino acid sequences of Pso2.

The plasmid p416-*PSO2::FLAG* constructs bearing mutation in the NLSs were individually transformed into the *S. cerevisiae pso2Δ* strain to determine changes in the intracellular localization and abundance of the ectopically expressed Flag-tagged wild-type Pso2 and its variants. A Western blot analysis was performed with an antibody specific to the FLAG-tag using the clarified nuclear lysates of unstressed cells (Fig. 5A). In multiple experiments, we observed a marked reduction in the amount of Pso2 that had mutations at either NLS1 or NLS2 motifs as compared to the wild-type (Fig. 1A). However, co-disruption of NLS1 and NLS2 motifs almost completely abolished the Pso2 nuclear import, indicating synergistic interactions between these motifs (Fig. 5B). Interestingly, while mutations in the MTS domain increased the translocation of Pso2 into the nucleus, import of its nuclease-deficient variant was reduced by 0.93-fold (Fig. 5C).

**Fig. 5.**
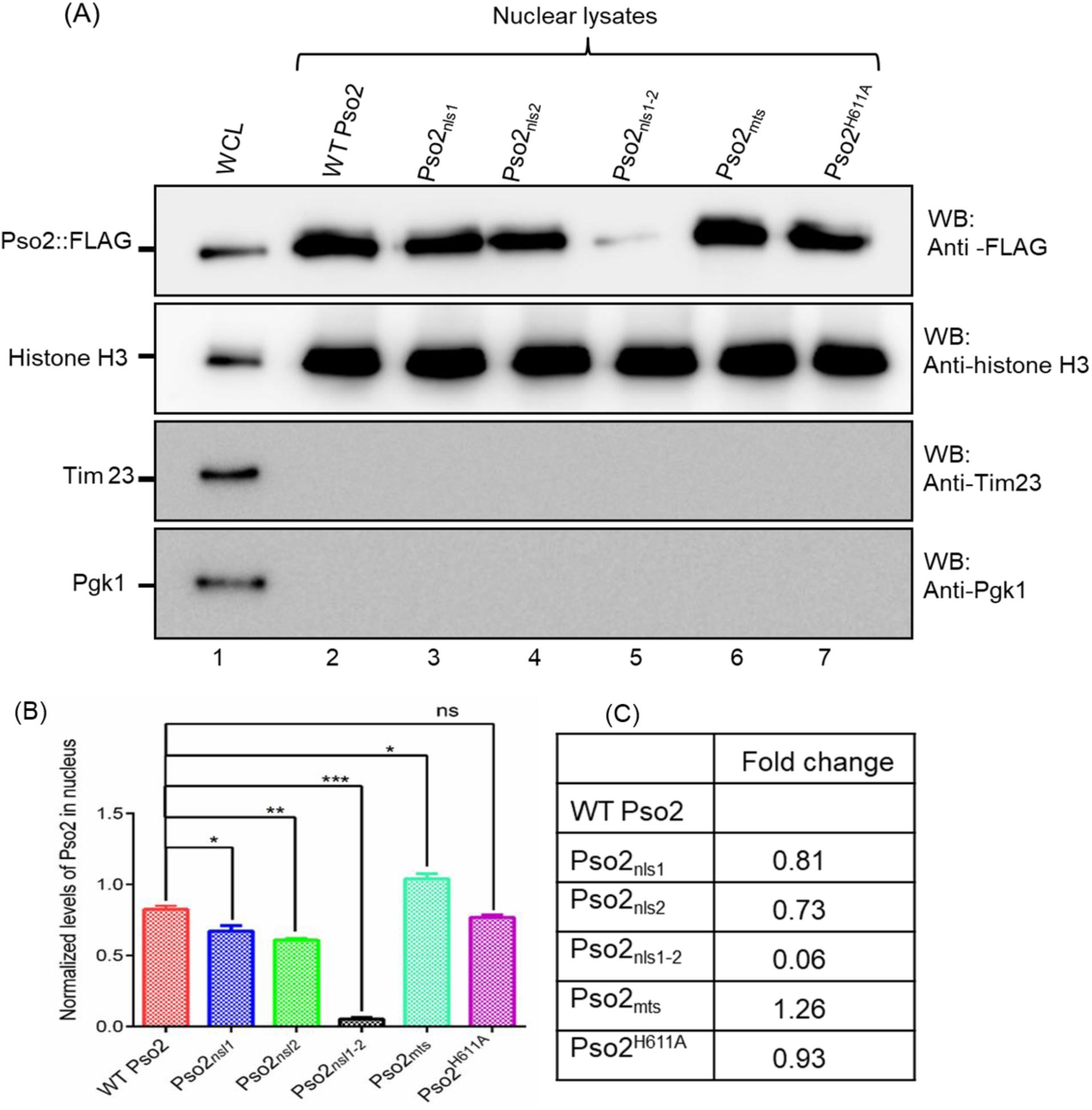
Either NLS1 or NLS2 of Pso2 is sufficient for its nuclear localization. (A) Western blot analysis of nuclear lysates. Upper panel, whole cell lysates (WCL) or nuclear lysates from *S. cerevisiae pso2Δ* cells expressing the wild-type Pso2-FLAG and its NLS and MTS variants were resolved by SDS-PAGE and probed analysis using antibody against the indicated protein. Lane 1 represents the whole-cell lysate (15 µg protein) from wild-type Pso2-FLAG; lanes 2-7, nuclear lysates (15 µg protein) from cells harbouring plasmids expressing Pso2-FLAG and its NLS, MTS and nuclease-inactive variants as indicated above each lane. While the blot in the upper panel was probed using anti-FLAG antibody, the blots depicted in the three lower panels were probed with anti-histone H3, anti-Tim23 and anti-Pgk1 antibodies, respectively. (B) Quantification of relative levels of signal intensity shown in (A). The data is expressed as the mean ± SEM from three independent experiments. Statistical comparisons were performed by unpaired Student’s t test (*p < 0.05, **p < 0.01, ***p < 0.001, ns, not significant). (C) The fold-change in the relative levels of Pso2 variants (compared with the wild-type) in the nucleus.

In individual immunoblotting experiments, we verified equal loading of nuclear proteins in each lane (histone H3) and excluded the possibility of cross contamination by mitochondrial (Tim23) and cytosolic (Pgk1) proteins, respectively (lower three panels of Fig. 5A). Tim23 and phosphoglycerate kinase have been widely used as marker proteins for assessing the purity of mitochondrial and cytosolic protein samples, respectively. Collectively, our data clearly demonstrate that either NLS1 or NLS2 was sufficient to drive nuclear import of Pso2 to almost similar levels, implying that the two NLS motifs are functionally redundant.

### While disruption of MTS motif impairs the mitochondrial translocation of Pso2, mutant NLS motifs enhance its localization to the mitochondria

After observing the robust effects of MTS and NLSs on the nuclear localization of Pso2, we utilized a similar approach to evaluate the impact of mutations on the translocation of Pso2 into the mitochondria. To this end, Western blot analysis was carried out on the clarified mitochondrial lysates of unstressed *pso2Δ* cells expressing the Flag-tagged wild-type Pso2 or variant species bearing mutations in the MTS or NLS motifs. Illustrative results in Fig. 6A show that while mutations in the NLS1 elevated translocation of Pso2 into the mitochondria, mutations in the NLS2 motif was significantly less robust (Fig. 6B). A possible explanation of this result would be that the MTS function is overwhelmed by NLS1 and, consequently, mutation in the latter rises its abundance in the mitochondria. Consistent with this notion, co-disruption of NLS1 and NLS2 motifs elevated the levels Pso2 in the mitochondria by 1.6-fold, as compared to the wild-type (Fig. 6C). Notably, point mutations of conserved residues in the MTS sequence markedly impaired Pso2 import into the mitochondria, although it was not abolished (Fig. 6B-C). There are several possible explanations. This could be explained by the existence of an alternative, although less efficient, mechanism of mitochondrial import or that point mutations may not fully disrupt the putative secondary structure acquired by MTS. Regardless, combined results endorse the view that positively charged residues in the MTS and NLS motifs play crucial roles in the nucleo-mitochondrial import of Pso2. Surprisingly, we found that the nuclease-deficient Pso2 variant was imported into the mitochondria with an efficiency similar to the wild-type (Fig. 6A-B). In control experiments, we verified loading of an equal amount of mitochondrial lysates per lane using anti-Tim23 antibody, and excluded the possibility of cross-contamination by cytoplasmic and nuclear fractions by immunoblotting against the indicated marker proteins (lower three panels in Fig. 6A). Taken together, these results support the notion that MTS and NLS motifs play an essential role in regulating the nucleo-mitochondrial translocation of Pso2.

**Fig. 6.**
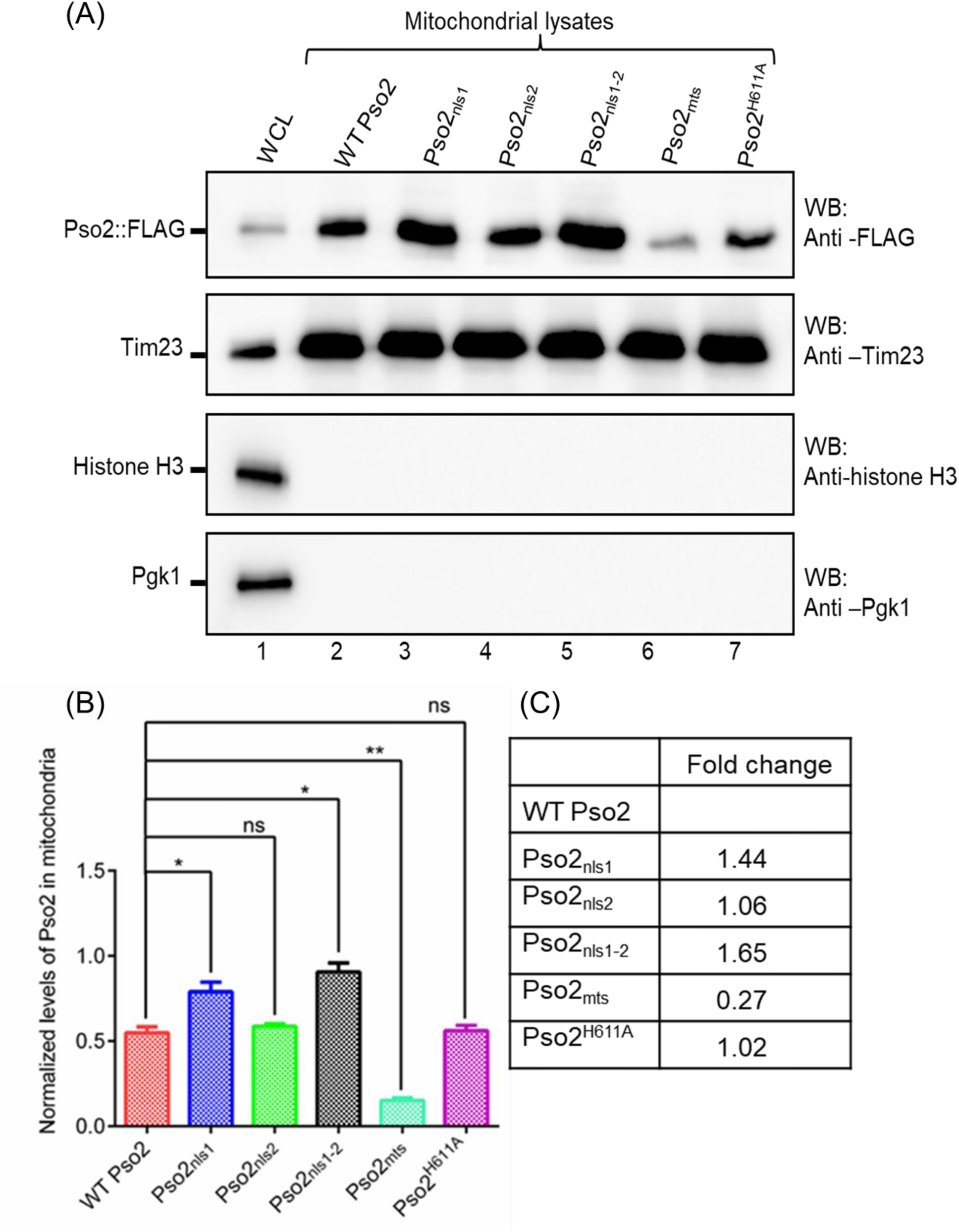
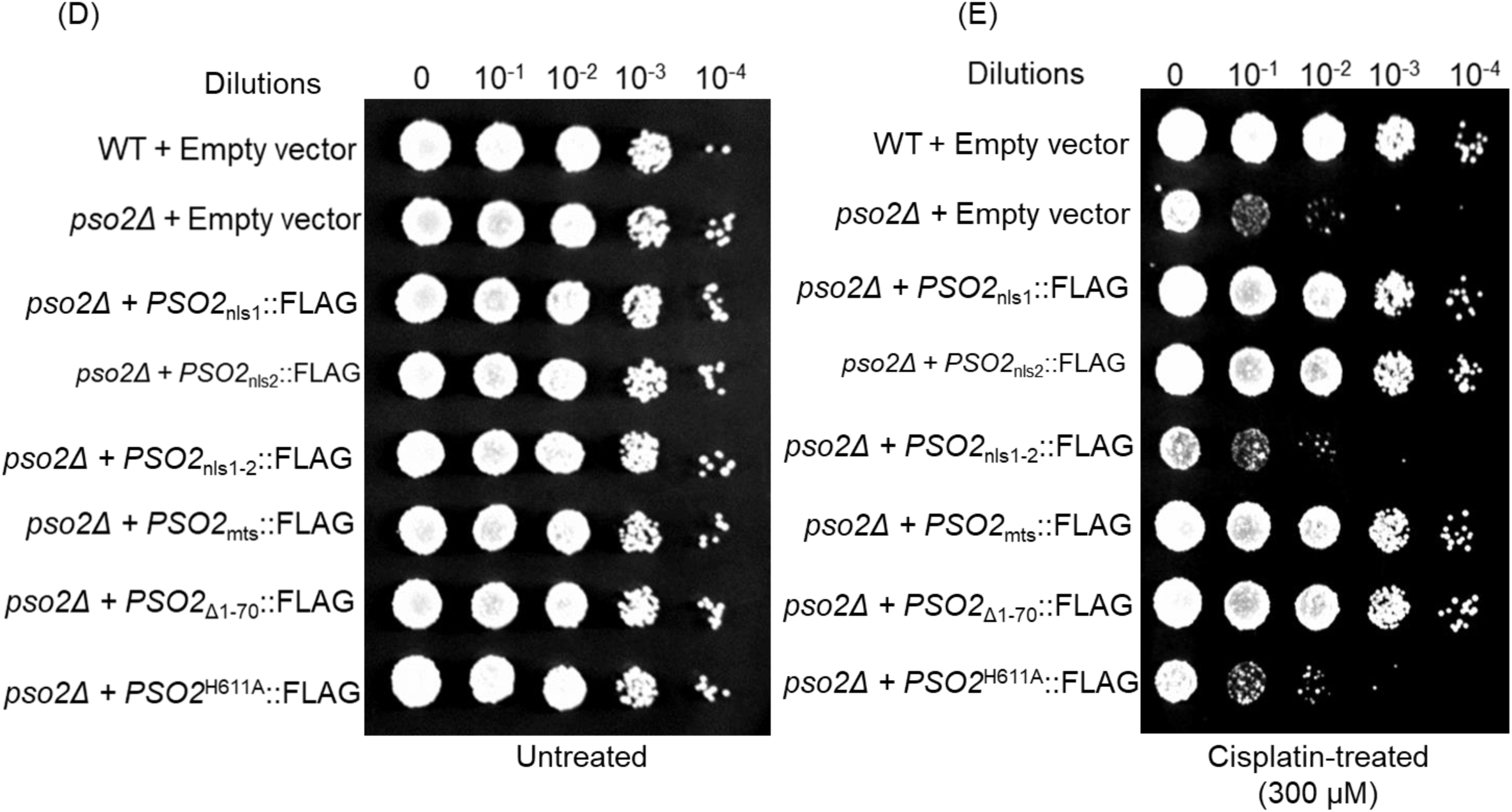
Point mutations within the putative MTS of Pso2 disrupt its mitochondrial localization. (A) Western blot analysis of mitochondrial lysates. Upper panel: whole-cell lysates (WCL) or mitochondrial lysates of *S. cerevisiae pso2Δ* cells expressing the wild-type Pso2-FLAG of its NLS and MTS variants were resolved by SDS-PAGE and probed with antibody against the indicated protein. Lane 1 represents the whole-cell lysate (15 µg) from wild-type Pso2-FLAG; lanes 2-7, mitochondrial lysates (15 µg) from cells harbouring plasmids expressing the wild-type Pso2-FLAG or its NLS, MTS and catalytically-inactive variants as indicated above each lane. While the blot in the upper panel was probed with anti-FLAG antibody, the blots depicted in the three lower panels were probed with anti-TIM23, anti-Ydj1 and anti-histone H3 antibodies, respectively. (B) Quantification of relative levels of signal intensity shown in (A). Statistical comparisons were performed by unpaired Student’s t test (*p < 0.05, **p < 0.01, ns, not significant). (C) The fold-change in the relative abundance of Pso2 variants in the mitochondria compared to that of the wild-type. (D) Growth of serially diluted untreated *S. cerevisiae* wild-type strain and its *pso2Δ* isogenic strains complemented with plasmids expressing the wild-type Pso2-FLAG, NLS or MTS variants of Pso2 on uracil-lacking SD medium. (E) As in panel (D), but the cells were treated with 300 μM cisplatin. The data shown is representative of three independent experiments.

### Growth phenotypes of cells expressing the wild-type and import-deficient Pso2 variants

To determine whether mutations in the MTS and NLS motifs of Pso2 affect cell growth in response to genotoxic stress, the ability of import-deficient variants of Pso2 to attenuate the sensitivity of *pso2Δ* cells to cisplatin was assessed. We surmised that the import-deficient Pso2 variants would be defective in ICL repair because of their inability to translocate into the mitochondria and nucleus. To investigate this, serially diluted *S. cerevisiae* wild-type cells carrying an empty vector, its isogenic *pso2Δ* cells bearing plasmids expressing the import-deficient Pso2 variants or nuclease-deficient mutant (pso2^H611A^) were spotted on a uracil-lacking SD medium, with or without cisplatin. As expected, the growth phenotypes of untreated cells expressing different Pso2 variants were very similar to that of the wild-type strain bearing an empty vector (Fig. 6D). On the other hand, while the wild-type strain bearing empty vector was resistant to cisplatin, *pso2Δ* cells bearing an empty vector and *pso2Δ* strain expressing Pso2 NLS_1-2_ variant were hypersensitive to cisplatin (Fig. 6E). However, the sensitivity of *pso2Δ* strain to cisplatin could be rescued by ectopic expression of Pso2 variants having point mutations in the NLS1 or NLS2 motifs. Surprisingly, the *pso2Δ* cells expressing either the Pso2_mts_ or pso2_Δ1-70_ variant (both MTS and NLS1 motifs deleted) phenocopied the wild-type strain bearing the empty vector. In agreement with a previous report (Rogers *et al*. 2020), the nuclease-deficient variant (Pso2^H611A^) failed to rescue *pso2Δ* cells from cisplatin-induced cell death (Fig. 6E), although significant amounts of mutant protein accumulates both in the nucleus and mitochondria (Fig. 5A-C, and 6A-C). These results reveal that (a) *pso2* NLS1/NLS2^-/-^ double variant scores as fully defective *in vivo*, (b) point mutations in NLSs, MTS or NLS1-MTS deleted species of *pso2* rescued the growth defect of *pso2Δ* strain, and (c) the nuclease-deficient pso2^H611A^ variant fails to recuse *pso2Δ* cells from cisplatin-induced cell death. We shall discuss these results further in the Discussion.

It is quite evident from the above data that mutational disruption of MTS and NLS motifs significantly decreased the levels of import-deficient Pso2 variants in the mitochondria and nucleus, respectively. However, the variations in their levels may be attributable to differential expression (or stability) of import-deficient Pso2 variants. To test this hypothesis, whole-cell lysates of *pso2Δ* cells expressing Flag-tagged wild-type Pso2 and its variants harbouring mutations in the MTS, NLS1 and NLS2 motifs were subjected to Western blot analysis and probed with antibodies specific for the FLAG-tag in Pso2. Illustrative data in Supplementary Fig. S1 show no significant differences between the levels of import-deficient of Pso2 variants and the wild-type in the whole-cell lysates. Taken together, these results are consistent with the notion that variation in the levels of import-deficient variants in the nucleus and mitochondria is not due to differences in their expression, but caused by mutations within the MTS and NLS motifs.

### Pso2 resides in the mitochondrial matrix compartment

Generally, mitochondrial proteins can localize to four distinct compartments: the outer membrane, inner membrane, inter membrane space, and protein-dense matrix (Fox, 2012). Three different approaches were employed to determine the location of Pso2 in the mitochondria. In this assay, Tom70, Tim23 and Mge1 served as markers of the outer membrane, inner membrane and matrix, respectively. In the first approach, isolated mitochondria from *S. cerevisiae pso2Δ* cells expressing the wild-type Pso2-FLAG were resuspended in an isotonic buffer and then treated with proteinase K (PK) to digest the protein(s) associated with the outer membrane. The samples were separated by SDS-PAGE, followed by Western blot analysis using anti-FLAG antibody. The results showed that whilst Tom70 was completely degraded by PK, Mge1 was not. Further, Pso2 remained resistant to PK digestion (Fig. 7A), indicating that it was not associated with the outer membrane. In the second approach, the protein associated with the inter membrane space, and matrix of intact mitochondria were extracted into the soluble fraction using an alkaline buffer (pH 11.5), and the samples were analysed as described above. The data showed that Pso2 was present in the supernatant fraction, indicating that it could be localized at the inter membrane space or the matrix (Fig. 7B). Finally, to further analyse whether Pso2 resides within the inter membrane space or matrix, mitoplasts were prepared by incubating intact mitochondria with hypotonic buffer. The resulting samples were analysed as described above. Notably, Pso2 was found in the mitochondrial matrix fraction (Fig. 7C). The combined results strongly support the notion that Pso2 resides in the internal matrix.

**Fig. 7.**
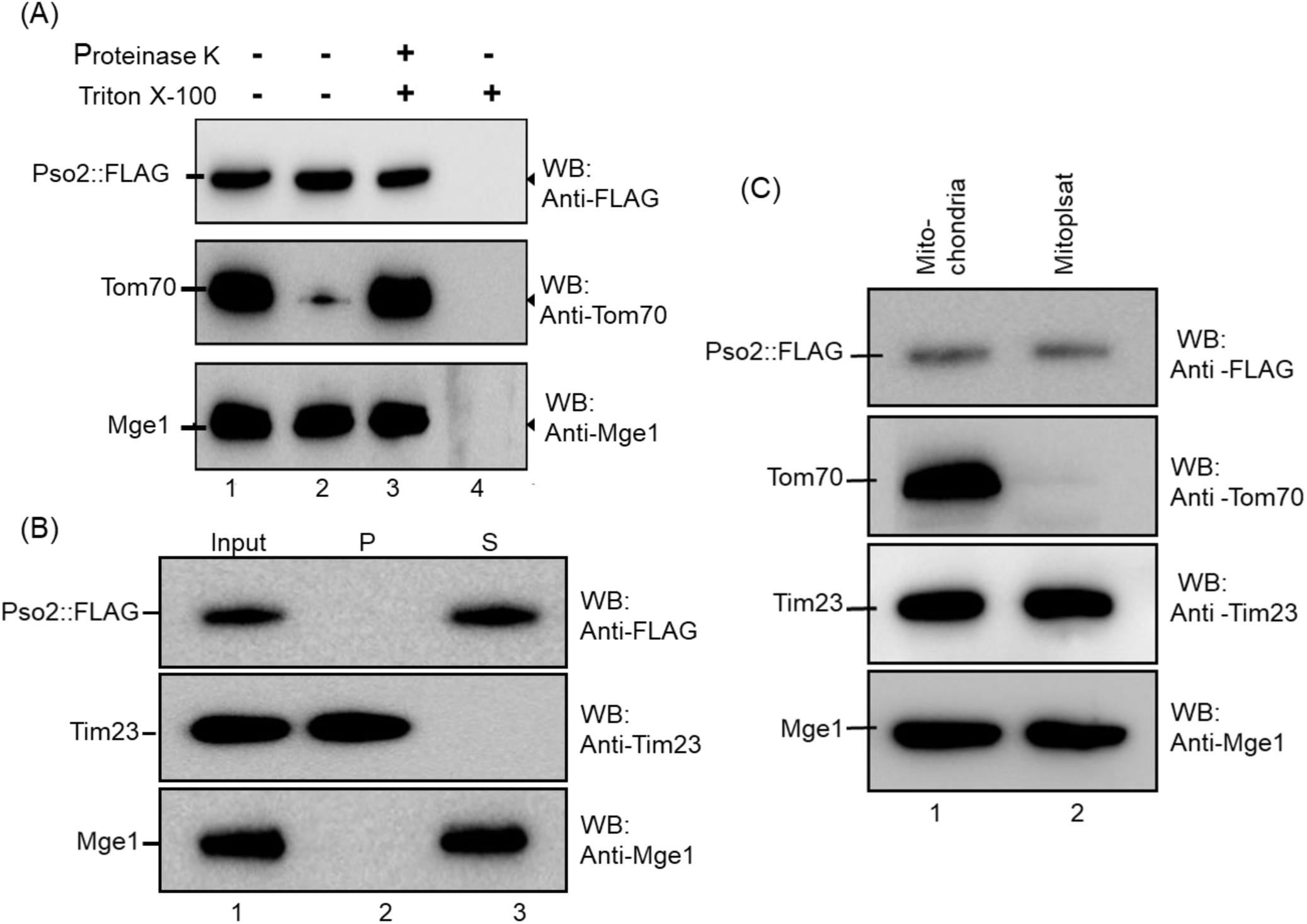
Pso2 resides in the mitochondrial matrix compartment. (A) Upper panel: Western blot analysis of samples using anti-FLAG antibody. Lane 1, untreated control. Lanes 2-4 represent mitochondrial lysate incubated in a buffer containing proteinase K, Triton X-100 and proteinase K with Triton X-100, respectively. (B) Upper panel: Western blot analysis of samples from alkaline extraction. Lane 1, untreated control. Lanes 2-3, samples from the pellet and supernatant fractions. (C) Upper panel, Western blot analysis of lysates of mitochondria (lane1) and mitoplast (lane 2). The blots in the lower two panels were probed with anti-Tom70, anti-Tim23 or anti-Mge1 antibodies as indicated.

### Point mutations in the MTS and NLS motifs do not blunt Pso2 exonuclease activity

Early biochemical studies provided evidence that Pso2 has an intrinsic 5′-to-3′ exonuclease and structure-specific endonuclease activity (Ma *et al*. 2002; Li *et al*. 2005; Tiefenbach and Junop, 2012). Recognizing that the MTS and NLS motifs are distributed along 30% of Pso2 polypeptide chain length, although noncontiguously, we sought to determine whether mutations in the NLS and MTS motifs affect its biological function. Hence, exonuclease activity was leveraged as a tool to assess the functional integrity of import-deficient variants of Pso2. To this end, wild-type Pso2 and its import-deficient variants were expressed as His-tagged proteins and purified individually from *E. coli* whole cell lysates via Ni^2+^-NTA affinity chromatography followed by Q Sepharose Fast Flow anion exchange chromatography (Fig. 8A-F). The overall yield of the import-deficient variants of Pso2 was similar to the wild-type, indicating no changes in their stability during expression and purification. The exonuclease activity of affinity purified proteins was analysed using radiolabelled ssDNA as the substrate. We titrated increasing concentrations of the wild-type Pso2 or its variants into the cleavage reactions containing 5’ ^32^P-labelled 60-mer ssDNA. The products of these reactions were electrophoresed through denaturing 15% polyacrylamide gels. The results from this analysis indicated that the amount of input substrate gradually declined in a similar fashion with increasing concentration of the wild-type and mutant species of Pso2 (Fig. 8G-K). In agreement with a previous report (Tiefenbach and Junop, 2012), the nuclease-deficient Pso2^H611A^ variant was completely devoid of exonucleolytic activity (Fig. 9L). Evaluation of individual cleavage efficiencies of import-deficient variants of Pso2 the wild-type enzyme revealed no significant differences (Fig. 8M). Taken together, these findings reinforce the notion that mutations in the MTS and NLS motifs do not affect the enzymatic activity of purified Pso2.

**Fig. 8.**
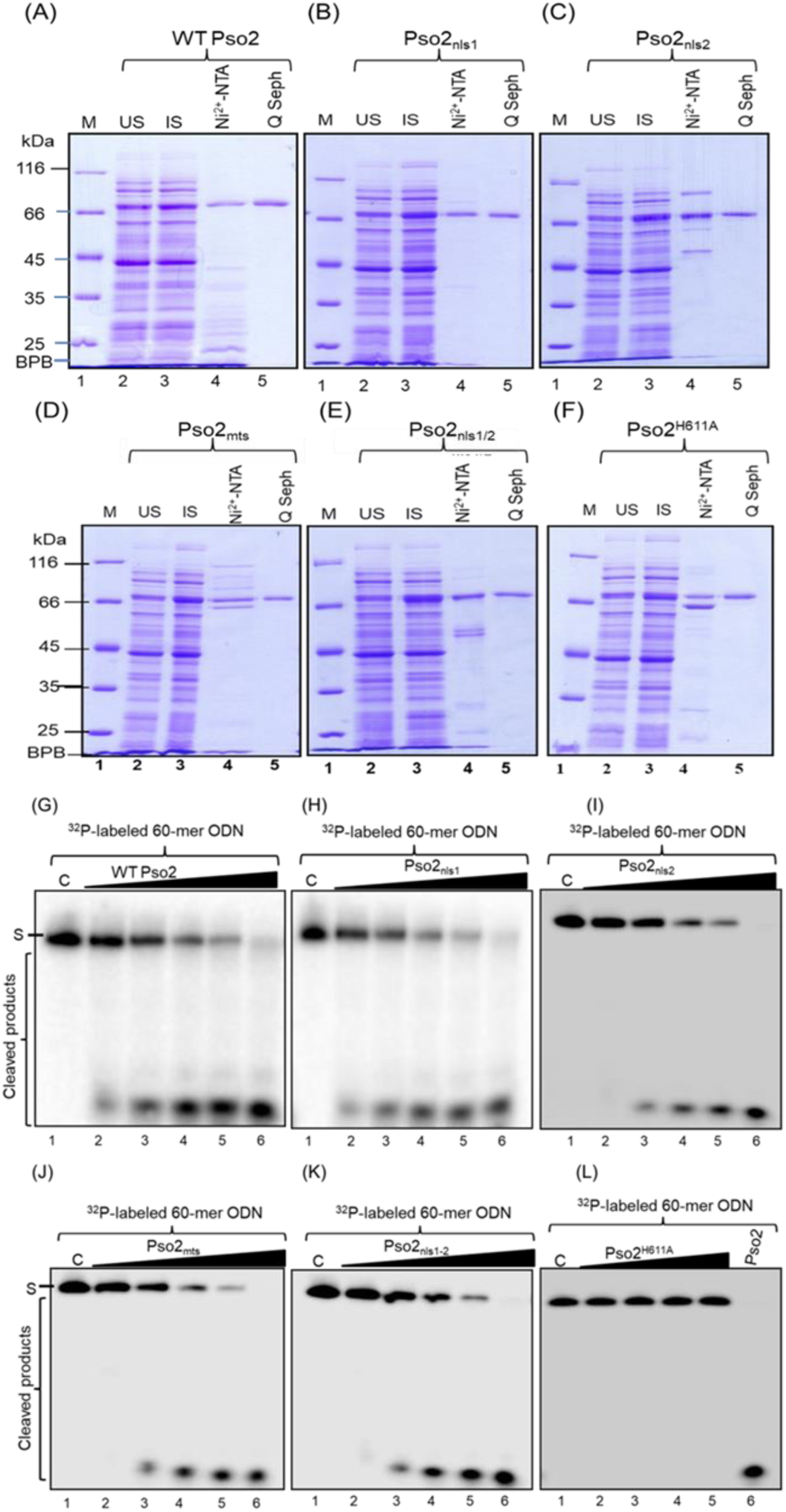
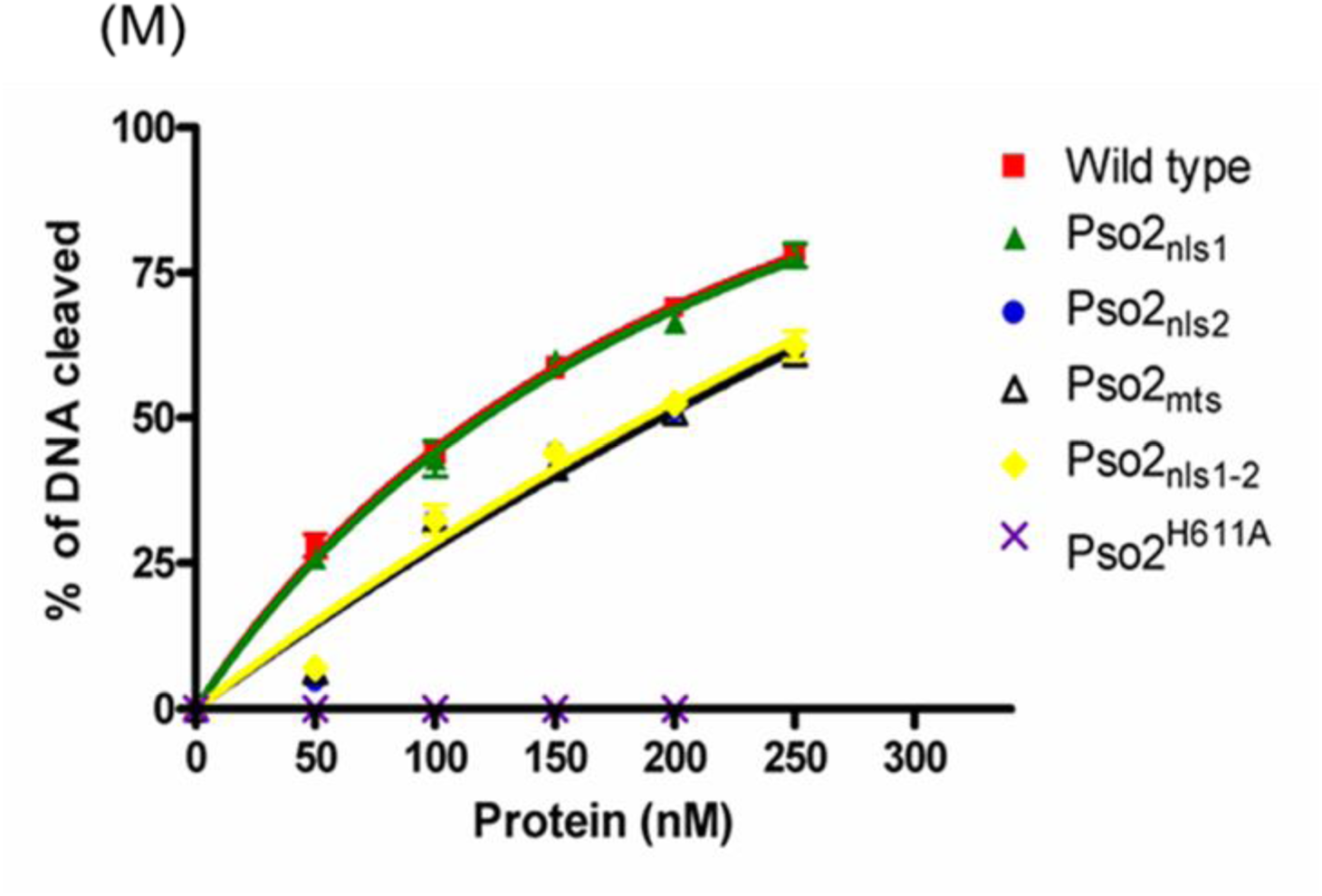
Mutations within the MTS and NLS sequences do not affect the exonuclease activity of Pso2. Panels A-F shows SDS-PAGE analysis of the expression and purification of wild-type Pso2 and its variant forms. In each panel, lane 1 shows molecular weight markers whose mass is given in kDa. Lane 2, uninduced whole cell lysate (15 □g); 3, induced whole cell lysate (18 □g); 4, elute from Ni^2+^-NTA column (8 □g); 5, elute from Q-Sepharose column (4 □g). Panels G-K show the 5’ exonuclease activity of the wild-type Pso2 and its variant forms; Pso2_nls1_, Pso2_nls2_, Pso2_mts_ and Pso2_nls1-2_ respectively. In each panel, lane C indicates no-enzyme control. Lanes 2-6 show the reactions carried out with the protein specified on the top of the gel at a concentrations of 50, 100, 150, 200 and 250 nM respectively. Panel L shows the 5’ exonuclease activity of Pso2^H611A^. Lane 1, control reaction in the absence of protein; lanes 2-5, assay performed with increasing concentration of Pso2^H611A^ at 50, 100, 150 and 200 nM, respectively. Lane 6, assay performed with 250 nM of the wild-type Pso2. S indicates the substrate. As the 60-mer ODN is labelled at the 5’end, the intermediate products are not visible in the gels. (M) Graph shows phosphor imaging quantification of exonuclease activity as a function of increasing concentration of Pso2. Each point represents the mean of the three independent experiments. The best-fit curve was obtained by subjecting the data sets to nonlinear regression analysis, in GraphPad Prism (version 5.0).

## DISCUSSION

One of the most consistent observations from published work is that mtDNA (compared to genomic DNA) is particularly susceptible to endogenous and exogenous oxidative/nitrosative stress, chemical carcinogens and chemotherapeutic drugs (Tretyakova *et al*. 2015; Kuhbacher and Duxin, 2020; Rong *et al*. 2021). The DNA lesions caused by the agents mentioned above can affect the integrity and functionality of the mitochondrial genome. Although canonical nuclear histones are lacking, growing evidence suggests that the mitochondrial nucleoid structure, and DNA compaction provide a certain degree of protection to mtDNA (Lee and Han, 2017). And yet, faced with the daunting task of combating oxidative DNA damage, cells have evolved mechanisms to deliver nuclear-encoded protein factors for BER into the mitochondria (Bannwarth *et al*. 2012; Liu *et al*. 2015). However, it is less clear whether the protein factors involved in different DNA repair pathways including ICL, NER, DSBR and NHEJ occur within mitochondria.

In this scenario, our findings contribute substantially to the understanding of the functions of NLS and MTS motifs of Pso2 in its subcellular distribution in ways that were previously unappreciated. Most notable is the demonstration that Pso2, a nucleus-encoded ICL repair protein, is a dual-localized nucleo-mitochondrial protein. Although a previous study reported that Pso2 localizes to the nucleus (Tkach *et al*. 2012), it remained unknown whether its nuclear import is autonomous or mediated by the NLS motif(s). In this regard, we found evidence that Pso2 contains a pair of NLSs and a conventional MTS at its N-terminus, which are essential for the nucleo-mitochondrial localization of Pso2, as mutations in these motifs severely compromised Pso2 function and localization.

Typically, nucleo-mitochondrial proteins harbour distinct targeting signals, which are recognized by both mitochondrial and nuclear import apparatus (Martin *et al*. 2015). However, Pso2 is unique in having NLS1 as a part of MTS, and that these two targeting signals exert their function in a competitive manner. Mutational disruption and biochemical analyses of NLSs provided robust evidence that either NLS1 or NLS2 is sufficient for the translocation of Pso2 into the nucleus, suggesting that the NLSs display redundancy with respect to their function. Thus, substitution of the conserved Lys/Arg residues by Ala in the NLSs abrogated nuclear localization of Pso2, thereby pointing to the importance of positively charged residues for its nuclear import. Similarly, replacement of two contiguous Arg and Lys residues in MTS by Ala abolished the mitochondrial import of Pso2. However, the exonuclease activity of Pso2 is not vulnerable to the adverse effects of mutations in the MTS and NLS motifs.

Two main types of approaches were used to establish the roles of protein targeting signals. First, the *S. cerevisiae pso2Δ* strain bearing Pso2::GFP and Pso2::FLAG constructs were utilized in CLSM and Western blot analyses, respectively. Consequently, the emitted GFP fluorescence obligatorily arises from the Pso2::GFP in the *S. cerevisiae pso2Δ* cells, thereby providing unambiguous evidence for the subcellular localization of Pso2. Although the fluorescence intensity of Pso2-GFP signal in mitochondria varied among cells treated with different genotoxic stresses, we consistently observed a marked increase in the GFP-Pso2 puncta density following treatment with MMS as compared to cisplatin or H_2_O_2_. However, the basis for the distinct effects of different genotoxic stress-inducing agents (MMS versus cisplatin and H_2_O_2_) is under investigation. Consistent with this, Western blot analysis of mitochondrial lysates confirmed the presence of higher levels of Pso2 in the mitochondria. Additional data hinted at potential crosstalk between MTS and NLS motifs with regard to the localization of Pso2 between the two spatially separated, DNA-containing organelles (Saki and Prakash, 2017).

Secondly, the ability of the wild-type and import-deficient variant species of Pso2 to attenuate the toxic effects of cisplatin was evaluated. While blocking Pso2 translocation to the nucleus caused severe growth defects in cisplatin-exposed *pso2Δ* cells, surprisingly, a similar effect was not evident when the import of Pso2 into the mitochondria was impaired. There exist several possible explanations for this. One possibility is that the repair of damaged mtDNA alone may not be sufficient to overcome cisplatin-inflicted damage to the nuclear genome and confer cisplatin-resistance. In a similar fashion, the lack of cisplatin-sensitive phenotype can be reconciled by the fact that small amounts of Pso2 (Fig. 6A) may be adequate for the repair of cisplatin-induced lesions in the smaller mitochondrial genome (compared to the nuclear genome) and restore cell growth and viability. Another consideration is that other sources, including protein factors or processes that are not associated with ICL repair may affect the ability of cells to overcome the toxic effects of cisplatin. It is also possible that a combination of these mechanisms might enable the subcellular distribution of Pso2. Further research is required to distinguish between these possibilities. The growth defect rescue observed in cisplatin-treated *pso2Δ* cells depends on Pso2’s enzymatic activity since the expression of a catalytic dead mutant does not rescue the cisplatin-sensitivity of *pso2Δ* cells. Regardless, reflecting on the physiological significance of Pso2, its human counterpart, *SNM1A*, is associated with cancer risk (Lee *et al*. 2016; Wang *et al*. 2016; Laporte *et al*. 2020); therefore, elucidation of the mechanism underlying the subcellular distribution of *SNM1A* would help in better understanding of its function in caner susceptibility.

The MTS and NLS motifs are necessary and sufficient for the import of Pso2 into the mitochondria and nucleus, respectively. Interestingly, our results are also consistent with the concept of an inverse relationship between the MTS and NLS motifs in regard to translocation of Pso2 to the nucleus and mitochondria. For instance, while mutations in NLSs abolished Pso2 import into the nucleus, at the same time, there was a marked increase in its enrichment in the mitochondrial matrix. In line with this, mutations in the MTS abrogated mitochondrial import of Pso2 and routed it to the nucleus. Several lines of evidence support a model in which DNA damage response regulates protein (re)localization and abundance at specified sites (Martin *et al*. 2015). Our results are consistent with earlier findings that the levels of *PSO2* mRNA dramatically increase following ICL-induced DNA damage (Wolter *et al*. 1996; Lambert *et al*. 2003). Our demonstration that Pso2 resides inside the mitochondria, endorse the notion that it is essential for DNA repair in this organelle. How is dual targeting of Pso2 to the mitochondria and nucleus regulated? It is known that certain types of *S. cerevisiae pso2* mutants can remove ICLs with normal kinetics, but they are repair-defective (Wilborn and Brendel, 1989; Grossmann *et al*. 2001; Ward *et al*. 2012); accordingly, we speculate that the defect in these mutants, at least, in part, could be due to their impaired nucleo-mitochondrial localization. Alternatively, or in addition, yet unidentified post-translational modification of Pso2 may regulate its intracellular distribution. Concordant with this idea, phosphorylation of Pso2 has been shown to modulate its exo- or endo-nucleolytic activity (Munari *et al*. 2014). Likewise, other types of post-translational modifications might influence the competition between MTS and NLS motifs; which in turn could influence translocation of Pso2 from the nucleus to mitochondria and *vice versa*. Regardless, our results are consistent with a model wherein individual NLSs are sufficient for nuclear import of Pso2, and failure to do so could cause defects in DNA repair.

Several studies have established that many proteins targeted to the mitochondrial matrix contain an MTS, consisting of 10-80 amino acid residue long domain, which is capable of folding into a positively charged amphipathic α-helix (Wiedemann and Pfanner, 2017). The MTSs are found in the proteins along their entire length, although they are located primarily at the N-terminal end. In contrast to internal MTSs, N-terminal MTSs are proteolytically removed by peptidases after import into the mitochondria (Neupert, 1997). Our finding that two species corresponding to the size of full-length (78 kDa) and truncated species (68 kDa) of Pso2 were seen in whole cell lysates suggest that the N-terminal MTS is cleaved off from the full-length protein after its import into the mitochondria.

Historically, *S. cerevisiae PSO2/SNM1* is the founding member of the *SNM1/PSO2* nuclease family of ICL repair factors (Henriques and Moustacchi 1980, 1981; Ruhland *et al*. 1981a; Ruhland et *al.* 1981b). Over the years, several orthologs of *PSO2* have been identified in vertebrates, including humans (Aravind 1999; Cattell *et al*. 2010; Baddock *et al*. 2020). The roles of *PSO2/SNM1* in different DNA damage repair pathways have been gleaned from biochemical and molecular genetic studies. These studies showed that *PSO2/SNM1* mutant cells are hypersensitive to irradiation and other DSB inducing agents, display ICL-induced chromosomal rearrangements, Pso2/Snm1 co-localise with known DNA repair factors, Pso2/Snm1 are recruited to the sites of DNA repair and digest DNA through an ICL *in vitro* (Baddock *et al*. 2020). Thus, a model emerges wherein failure in the nucleo-mitochondrial localization of an otherwise catalytically active *SNM1* may contribute to one or more of the above mentioned defects/phenotypes. Regardless, to our knowledge, this is the first study implicating that protein localization signals play a vital role in loss-of-function Pso2 phenotypes *in vivo*. Therefore, by extension, it will be interesting to determine whether the vertebrate orthologs recapitulate the subcellular localization patterns of *S. cerevisiae* Pso2.

## REAGENT AND DATA AVAILABILITY

The strains and detailed protocols will be provided upon request. We affirm that all data necessary for confirming the conclusions of the article are present within the article, Figures and tables.

## ACKNOWLEDGEMENTS

We would to express our sincere thanks to Murray Junop and Patrick D’Silva for providing plasmids pDEST14-*PSO2* and pDEST14-*pso2*^H611A^, and pRS413-MTS::mCherry, respectively. We appreciate Patrick D’Silva for his gift of polyclonal antibodies against Tim23, Tom70, Mge1, and Ydj1, and Manoj Thakur for his assistance with statistical analysis.

## CONTRIBUTION STATEMENT

Conceptualization: S.C.S. and K.M.; Methodology and investigation: S.C.S; Analysis of data: S.C.S. and K.M.; Validation: S.C.S; Writing: K.M. with feedback from S.C.S. Both authors confirm that they have read and approved the final manuscript.

